# SHP2 Inhibition Abrogates Adaptive Resistance to KRAS^G12C^-Inhibition and Remodels the Tumor Microenvironment of *KRAS*-Mutant Tumors

**DOI:** 10.1101/2020.05.30.125138

**Authors:** Carmine Fedele, Shuai Li, Kai Wen Teng, Connor Foster, David Peng, Hao Ran, Paolo Mita, Mitchell Geer, Takamitsu Hattori, Akiko Koide, Yubao Wang, Kwan H. Tang, Joshua Leinwand, Wei Wang, Brian Diskin, Jiehui Deng, Ting Chen, Igor Dolgalev, Ugur Ozerdem, George Miller, Shohei Koide, Kwok-Kin Wong, Benjamin G. Neel

## Abstract

*KRAS* is the most frequently mutated oncogene in human cancer, and KRAS inhibition has been a longtime therapeutic goal. Recently, inhibitors (G12C-Is) that bind KRAS^G12C^-GDP and react with Cys-12 were developed. Using new affinity reagents to monitor KRAS^G12C^ activation and inhibitor engagement, we found that, reflecting its action upstream of SOS1/2, SHP2 inhibitors (SHP2-Is) increased KRAS-GDP occupancy, enhancing G12C-I efficacy. SHP2-Is abrogated feedback signaling by multiple RTKs and blocked adaptive resistance to G12C-Is *in vitro*, in xenografts, and in syngeneic *KRAS*^G12C^*-*mutant pancreatic ductal adenocarcinoma (PDAC) and non-small cell lung cancer (NSCLC) models. Biochemical analysis revealed enhanced suppression of ERK-, MYC-, anti-apoptotic-, and cell-cycle genes, and increased pro-apoptotic gene expression in tumors from combination-treated mice. SHP2-I/G12C-I also evoked favorable changes in the immune microenvironment, decreasing myeloid suppressor cells, increasing CD8+ T cells, and sensitizing tumors to PD-1 blockade. Experiments using cells expressing inhibitor-resistant SHP2 showed that SHP2 inhibition in PDAC cells is required for tumor regression and remodeling of the immune microenvironment, but also revealed direct inhibitory effects on angiogenesis resulting in decreased tumor vascularity. Our results demonstrate that SHP2-I/G12C-I combinations confer a substantial survival benefit in PDAC and NSCLC and identify additional combination strategies for enhancing the efficacy of G12C-Is.

## INTRODUCTION

The RAS/ERK mitogen-activated protein kinase (MAPK) cascade is among the most frequently affected pathways in human cancer (1-3). Mutations in genes encoding pathway components, including RTKs, SHP2, NF1, RAS or RAF family members, or MEK1/2 cause inappropriate pathway activation and promote oncogenesis. *RAS* (*KRAS, HRAS, NRAS*) mutations are found in ∼20% of all human neoplasms, contributing to ∼3.4 million new cases/year worldwide (4). *KRAS* is the most often altered *RAS* isoform in solid tumors: nearly all pancreatic ductal adenocarcinomas (PDACs), ∼50% of colorectal carcinomas (CRCs), and 25%-30% of non-small cell lung cancers (NSCLCs) express mutant KRAS. These mutations almost always (∼95%) affect codons 12, 13, or 61, markedly increase the RAS-GTP/RAS-GDP ratio, and activate effectors inappropriately (5, 6).

Mutant RAS was once viewed as impervious to GAP-stimulated or intrinsic hydrolysis, “locked” in the GTP state, and “undruggable.” More recent structural/biochemical analyses revealed subtle, but critical, differences in intrinsic and residual GAP-catalyzed GTPase activity, intrinsic and SOS-stimulated exchange, and effector-binding between RAS mutants (7-12). Some oncogenic mutants, notably KRAS^G12C^ (G12C) and to a lesser extent, KRAS^G12D^ (G12D), retain significant intrinsic GTPase activity. GTP hydrolysis in G12C is refractory to (and might be inhibited by) RAS-GAP; G12D, G12A, G12R, G12V, and Q61L/Q61H retain some GAP-responsiveness, and thus could undergo at least some KRAS-GTP hydrolysis in cells.

Recent successes in developing clinical grade G12C inhibitors (G12C-Is) illustrate why these details are so important (13-17). Such agents bind an evanescent pocket in KRAS-GDP, positioning a reactive group to couple to the mutant cysteine. Four are now in Phase I trials (AMG510, MRTX849, JNJ74699157, LY3499446) (18), and there are initial reports of efficacy in NSCLC patients with *KRAS*^*G12C*^ mutations (19-22). For such drugs to engage G12C, hydrolysis sufficient to generate RAS-GDP must occur. As G12C is GAP-refractory, only agents that inhibit exchange (as opposed to enhancing GAPs) can increase occupancy of the KRAS^G12C^-GDP state and thereby enhance the ability of G12C-Is to couple to mutant KRAS. Consequently, SOS1/2 can effectively be viewed as competitors of G12C-Is (and vice versa).

SHP2, encoded by *PTPN11*, comprises two SH2 domains (N-SH2, C-SH2), a catalytic (PTP) domain, and a C-terminal domain with two tyrosine residues that, when phosphorylated, bind GRB2. In its “closed” (inactive) state, the N-SH2 occludes the SHP2 PTP domain, blocking substrate access, while the PTP domain contorts the N-SH2, rendering it unable to bind phosphotyrosyl (pY-) peptides(23-25). Conversely, pY-peptide binding drives SHP2 to the “open” state. Most N-SH2 ligands belong to bis-pY motifs in RTKs, cytokine receptors, “scaffolding adapters” (e.g., GAB, IRS, FRS proteins), or immune checkpoint receptors. This elegant “molecular switch” ensures SHP2 activation in response to appropriate signals at proper cellular locales, and has been exploited to develop potent, selective, orally available allosteric SHP2-Is (25-30). These agents bind a previously unrecognized pocket in “closed” SHP2, acting as “molecular glue” to impede the N-SH2/loop/C-SH2 movements needed for activation (25, 31, 32). Four such drugs are also in Phase I trials (TNO155, RMC4630, JAB3068, RLY1971), and an initial efficacy signal for RMC4630 in *KRAS*-mutant NSCLC was reported recently (33). SHP2 is required for full activation of RAS and the RAS/ERK cascade, but whether SHP2 regulates RAS-exchange or RAS-GAP had been unclear. Recently, several groups, including ours, provided strong evidence that SHP2 acts upstream of SOS1/2 to regulate exchange; consequently, SHP2-Is abrogate adaptive resistance to BRAF- or MEK-inhibitors (28, 34-37). Recent reports (and our unpublished observations; see Results) show that *KRAS*^*G12C*^-mutant cancer cell lines treated with G12C-Is also develop adaptive resistance (22, 38-41). These studies reported that adaptive response to G12C-Is could be minimized by combining G12C-I with RTK or SHP2 inhibitors (22, 38, 40, 41). Some of these findings were validated in human cell-derived (CDXs) or patient-derived xenografts (PDXs) (39).

Tumors are not, however, mere collections of neoplastic cells. Rather, they resemble defective “mini-organs” with complex interactions between cancer cells and cells of the tumor microenvironment (TME), which includes resident and infiltrating immune, mesenchymal, and endothelial cells (42, 43). G12C-Is are mutant-specific and thus have direct effects only on *KRAS*^*G12C*^-mutant tumor cells. Nevertheless, they could modulate the TME by altering tumor cell production of growth factors, cytokines, and chemokines (20). Most other targeted agents, including SHP2-Is, can affect RAS/ERK signaling in normal, as well as neoplastic cells. SHP2 also has effects on parallel pathways (e.g., JAK/STAT signaling), and is implicated as an effector of inhibitory signaling by PD1 and some other immune checkpoint receptors (44-46).

A sophisticated understanding of cancer therapeutics requires delineation of tumor cell-autonomous and -non-autonomous actions. Here, we report the effects of G12C-I, SHP2-I, and G12C-I/SHP2-I combinations in syngeneic *KRAS*^*G12C*^-mutant PDAC and NSCLC models. We find that G12C-I/SHP2-I efficacy derives from effects on tumor cells and cells in the TME, and reveal direct anti-angiogenic effects of SHP2-Is.

## RESULTS

### SHP2 inhibition enhances KRAS-G12C inhibitor effects in PDAC and NSCLC cell lines

Allosteric SHP2 inhibitors (e.g., SHP099) reduce the activation of KRAS mutants that retain significant intrinsic GTPase activity (“cycling mutants”), most notably, KRAS^G12C^, and to a lesser extent, KRAS^G12D^ and KRAS^G12V^ (hereafter G12C, G12D, 12V) in cancer cell lines and reconstituted “*RAS*-less MEFs” (28, 34-37). As G12C is impervious to RAS-GAPs (8), these and other data established that SHP2 acts upstream of SOS1/2. We hypothesized that SHP2 inhibition, by decreasing SOS1/2 activity, would increase occupancy of the KRAS^G12C^-GDP state, thereby potentiating the effect of G12C-Is. To test this hypothesis, we assessed the effects of SHP099, the G12C-I ARS1620 (ARS), SHP099/ARS, or Vehicle (DMSO) control on the proliferation of *RAS*-less MEFs” reconstituted with KRAS mutants (Figure 1A). Consistent with our previous results (35), SHP099 inhibited WT-reconstituted MEFs, whereas ARS had no effect. By contrast, SHP099 and ARS each inhibited G12C-MEFs to some extent, but SHP099/ARS had far greater efficacy. Neither SHP099 nor ARS alone or in combination significantly impaired the proliferation of G12D-or Q61R-reconstituted cells. There was no difference in ARS-induced adaptive resistance in *Kras*^*wt*^/*KRAS*^*G12C*^ and *Kras*^*-/-*^/*KRAS*^*G12C*^ MEFs (Supplemental Figure 1A), suggesting a more important role for mutant KRAS in promoting adaptive resistance (see Discussion).

**Figure 1:**
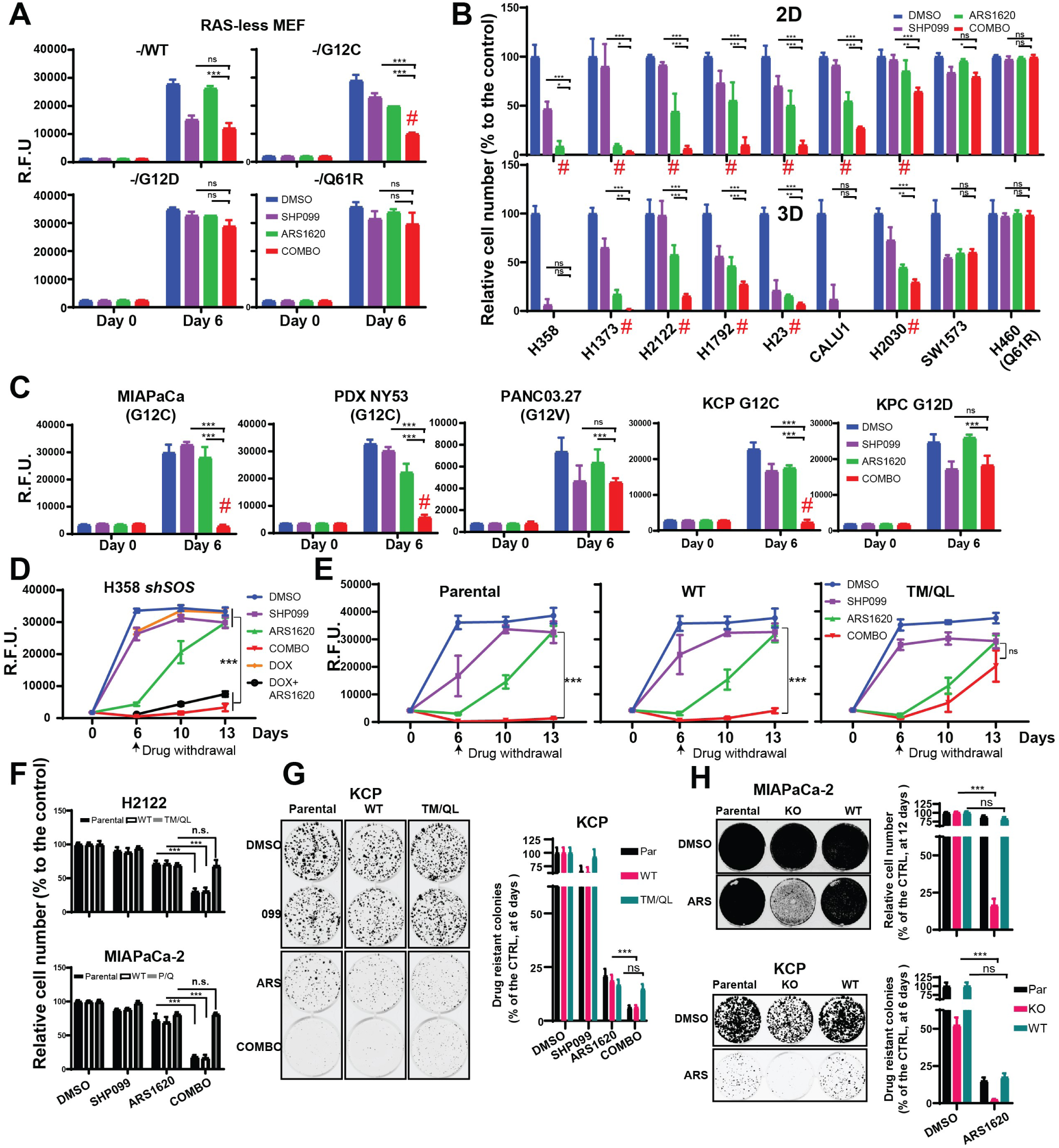
SHP2 inhibition enhances KRAS-G12C inhibitor effects in PDAC and NSCLC cell lines. **A-C**, Reconstituted RAS-less MEF (A), human NSCLC (B) and human and mouse PDAC (C) cell lines were treated with DMSO, SHP099, ARS1620, or both drugs (Combo). Cell viability, by PrestoBlue assay, was assessed at 6 days. **D**, Cell viability assays on H358 NSCLC cells expressing DOX-inducible sh*SOS1* (sh*SOS*) treated with DMSO, SHP099, ARS1620, or Combo for the indicated times. All drugs were withdrawn after 6 day of treatment and regrowth was tested at Days 10 and 13.**E**, Cell viability assay on H358 NSCLC cells expressing either SHP099-resistant *PTPN11* mutant (T253M/Q257L) or wild-type *PTPN11* (WT) treated with DMSO, SHP099, ARS1620, or Combo for the indicated times. All drugs were withdrawn after 6 days of treatment, and regrowth was tested at Days 10 and 13. **F**, PrestoBlue assay on H2122 and MIAPaCa-2 cells expressing SHP099-resistant *PTPN11* (T253M/Q257L or P491Q respectively) or wild-type P*TPN11* (WT) after 6 days of treatment. **G**, Colony formation assays (6 days) on KCP 1203 cells expressing *PTPN1*1-T253M/Q257L or wild-type *PTPN11* (WT). **H**, Colony formation assay (12 days) on *PTPN11*-KO or WT-*PTPN11*-reconstituted MIAPaCa-2 (12 days, top) and KCP 1203 (6 days, bottom) cells. Representative results are shown from a minimum of three biological replicates per condition. For all experiments, drug doses were: SHP099 10 μm/L, ARS1620 10 μm/l, Combo = SHP099 10 μm/l + ARS1620 10 μm/l. Data represent mean ± SD; *P< 0.05; **P < 0.01; ***P < 0.001, one-way ANOVA followed by Tukey’s multiple comparison test. Red symbols indicate synergistic interaction between the two drugs by BLISS independent analysis

Next, we tested a panel of *KRAS*^*G12C*^-mutant NSCLC lines cultured in monolayer (2D) or spheroid (3D) conditions; previous studies showed that 3D cultures are more dependent on the RAS/ERK pathway and thus are more sensitive to pathway inhibition (15, 16). Single agent SHP099 or ARS inhibited 2D-proliferation to varying extents, and, as expected, generally had greater effects on 3D proliferation (Figure 1B, Supplemental Figure 1B). Again, however, SHP099/ARS was far more effective than either agent alone on nearly all *KRAS*^*G12C*^ line. In most cases, the anti-proliferative effect of combining the inhibitors was synergistic (Figure 1B, red symbols, Table S1). By contrast, the *KRAS*^*G12C*^ line SW1573 failed to respond; these cells express *PIK3CA*^*K111E*^, a known gain-of-function allele, which might render them KRAS mutant-independent. As expected H460 cells, which harbor *KRAS*^*Q61R*^, were unresponsive to either single agent or the combination (Figure 1B). SHP099/ARS also showed greater ability than SHP099 or ARS alone to inhibit the proliferation of *KRAS*^*G12C*^-mutant MIAPaCa-2 PDAC cells, cells derived from a *KRAS*^*G12C*^-mutant patient-derived PDAC xenograft (PDX-NY53), and mouse PDAC cells (KCP) engineered from a *KRas*^*G12D*^*/Tp53*^*R172H*^ cell line (KPC) to have a single *Kras*^*G12C*^ allele (Figure 1C, Supplemental Figure 1B-C) and two inactive *Kras* alleles (Supplemental Figure 1D-I). Proliferation of KRAS^G12V^-expressing PANC03.27 cells and parental KPC cells was inhibited to some extent by SHP099, but as expected, ARS had no effect alone nor any additional effect when combined with SHP099 (Figure 1C, Supplemental Figure 1B-C). Notably, SHP099/ARS enhanced cell death (measured at 48h of treatment), compared with either single agent (Supplemental Figure 1J), most likely explaining its increased anti-proliferative actions. Some newer G12C-Is are more potent than ARS (20, 22). Nevertheless, SHP099 also enhanced the effects of the clinical grade inhibitor AMG510 (Supplemental Figure 1K).

If SHP2 inhibition potentiates G12C-I action by diminishing SOS activation, then *SOS* down-regulation should phenocopy the effects of SHP099. To test this possibility, we generated H358 cells expressing doxycycline (DOX)-inducible *shSOS1*. Indeed, ARS inhibition and *SOS1* shRNA expression had similar effects to SHP099/ARS treatment (Figure 1D). Expression of *PTPN11*^*T253M/Q257L*^, a mutant predicted to lack SHP099 binding (26), eliminated the effects of SHP099 in combination-treated H358 and KCP cells (Figure 1E-G). Similarly, another drug-resistant mutant, *PTPN11*^*P491Q*^, rescued the effects of SHP099/ARS on MIAPaCa-2 cells (Figure 1F). Moreover, ARS had similar effects on *PTPN11* knock-out (KO)-MIAPaCa-2 and -KCP cells (generated by CRISPR/Cas9) as did SHP099/ARS on parental or *PTPN11*-reconstituted cells (Figure 1H, Supplemental Figure 1L). Taken together, these findings establish that SHP099 and ARS are “on-target” and that SHP2 inhibition improves the effect of G12C-Is in multiple *KRAS*^*G12C*^-mutant cancer cell lines, arising from two distinct tissues.

To facilitate more direct assessment of G12C-I action, we used two novel affinity reagents for “pull-down” experiments. First, we employed a recently developed “monobody” (12C/V-MB) that selectively binds KRAS^G12C^-GTP but not KRAS^G12C^-GDP, and to a lesser extent, KRAS^G12V^ (47). We also used phage display to isolate a synthetic Fab (12C-ARS-Ab) that specifically recognizes ARS-adducted G12C with high affinity (Figs. 2A, Supplemental Figure 2A and data not shown; see Methods). Used in concert, 12C/V-MB and 12C-ARS-Ab affinity purifications, followed by RAS immunoblotting (hereafter, “pull down” assays), provide reciprocal information on the amount of KRAS^G12C^-GTP and KRAS^G12C^-ARS complexes.

**Figure 2:**
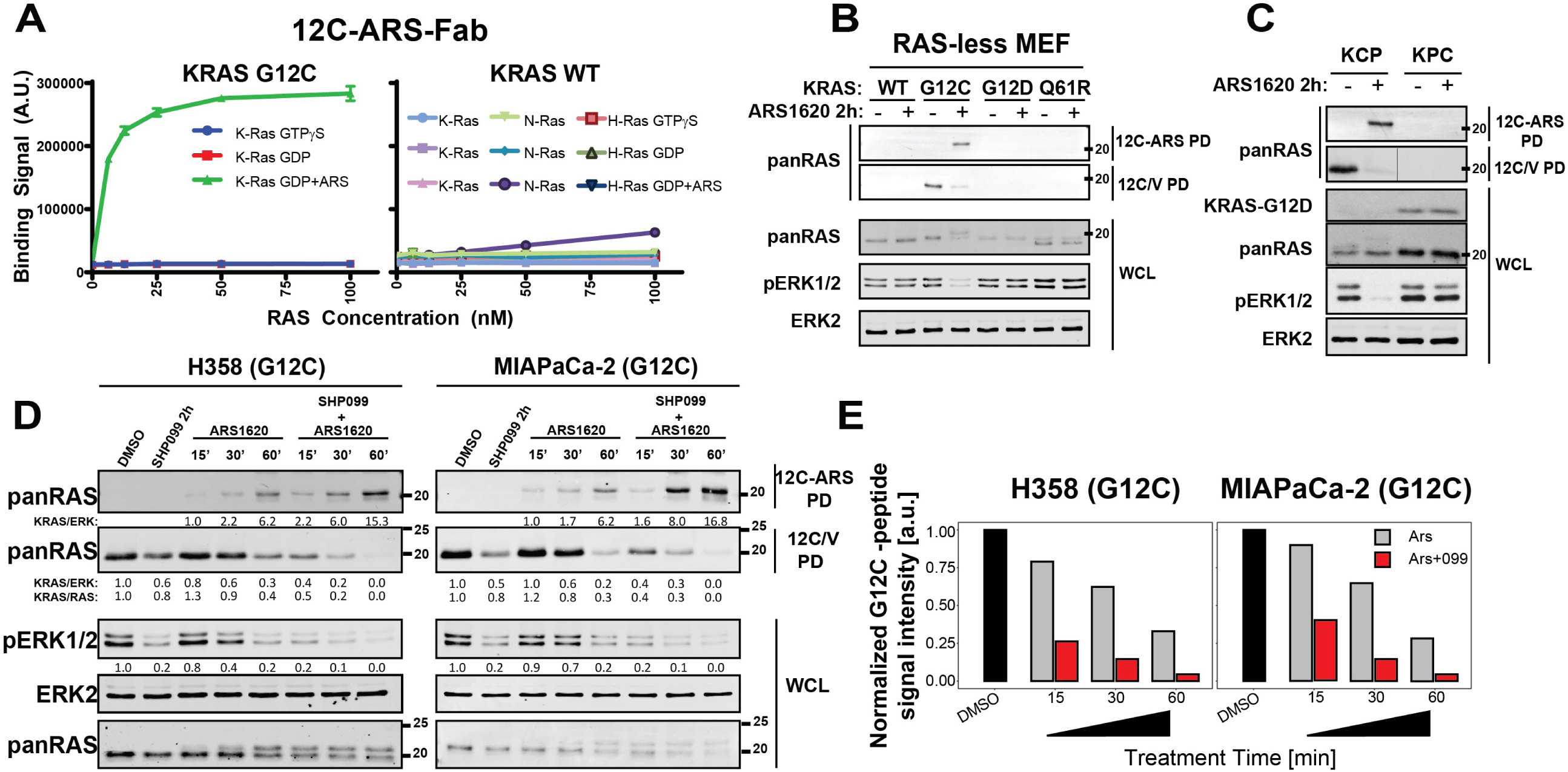
SHP099 increases KRAS-G12C-ARS1620 adducts. **A**, 12C-ARS Fab binding to KRAS-G12C (left) or WT RAS isoforms (right) with/without ARS and GTPγS or GDP. **B-C**, Immunoblots of whole cell lysates and 12C/V MB- or 12C-ARS Fab-pull-downs (PD) from RAS-less MEFs reconstituted with the indicated RAS mutants (B) and KCP cells (C), treated in the presence or absence of ARS1620. **D**, Immunoblots of whole cell lysates and 12C/V MB- or 12C-ARS Fab pull-downs (PD) from H358 and MIAPaCa-2 cells, treated as indicated. **E**, ARS-adduct formation in samples from C, quantified by LC/MS-MS. ARS1620 and SHP099 concentrations were 10 μm/l in all panels.

We validated these new reagents by testing lysates from MEFs reconstituted with WT-*KRAS* or various *KRAS* mutants and from KCP and KPC mouse PDAC cells. Cells were treated with or without ARS (2h), and lysates were subjected to “pull down” (PD) assays. ARS treatment lowered KRAS^G12C^-GTP levels, as indicated by decreased RAS signals in the 12C/V-MB PDs from G12C-expressing MEFs or KCP cells but not from the other lines. Conversely, there was an increase in RAS signal in 12C-ARS-Ab PDs from RAS-less MEFs reconstituted with KRAS^G12C^, but not other *KRAS* mutants, as well as from KCP, but not KPC, cells (Figure 2B-C).

These reagents provide a direct assessment of G12C-I action and thus enable direct assessment of how SHP099 potentiates the effects of ARS. To this end, we treated H358 and MIAPaCa-2 cells with ARS alone, SHP099 alone, or SHP099/ARS for various times and performed PD assays. The reciprocal recovery of KRAS in 12C/V-MB and 12C-ARS Fab PDS from ARS-treated cell lysates demonstrated time-dependent formation of ARS-adducts. Pre-treatment with SHP099 accelerated ARS-adduct formation across the time course: for example, complete engagement of KRAS^G12^ by ARS was seen by 1h in SHP099/ARS-treated cells, compared with ∼70 % engagement in cells treated with G12C-I alone (Figure 2D). These events were paralleled by more efficient pERK inhibition and slower mobility (in SDS-PAGE) of mutant KRAS upon SHP099/ARS-treatment (Figure 2D and Supplemental Figure 2A-B). Similar potentiation of G12C-I engagement by SHP099 was observed by using 12C/V-MB PDs on lysates from AMG510- and AMG510/SHP099-treated MIAPaCa-2 or H358 cells (Supplemental Figure 2C). We validated these findings by using the current gold standard MS assay (15), which measures a decrease in C12– containing peptide relative to isotopic standards (G12C peptide: LVVVGACGVGK; KRAS/NRAS normalization peptide: SYGIPFIETSAK, both spiked into lysates) in tryptic digests of ARS- or SHP099-treated cell lysates (Figure 2E). These results, along with the known biochemical properties of G12C (retained intrinsic GTPase, GAP-non-responsive), provide further, incontrovertible evidence that SHP2 inhibitors impede RAS-GEF action.

### SHP099 abrogates adaptive resistance to G12C-Is *in vitro*

MEK-I treatment of *KRAS*-mutant tumors fails, at least in part due to induction of genes encoding multiple RTKs and/or their ligands, which differ between tumors even of a single histotype (48-51). We (35) and others (28, 34-37, 52, 53) reported that SHP2 inhibitors, by blocking RAS activation evoked by MEK-I-induced RTKs/RTK ligands, could block adaptive resistance to MEK-Is and that SHP2-I/MEK-I combinations synergistically inhibited the proliferation of multiple *KRAS*-mutant cancer models. We analyzed RTK and RTK ligand gene expression in ARS-treated MIAPaCa-2 and H358 cells by qRT-PCR (Figure 3A). Several—but different--RTKs were induced by G12C-I treatment, including *EGFR, FGFR3, IGFR1, MET, VEGFR1* and *PDGFRA/B* in MIAPaCa-2 cells, *ERBB2/3, FGFR2/3* and *PDGFRA/B* in H358 cells. The same lines variably induced *EGF, FGF2, PDGFB, PDGFC, PDGFD* and/or *VEGFA/B* RNA. Consequently, it is difficult, if not impossible, to design an efficient combination therapy with G12C-Is by targeting RTKs directly. Notably, the RTK/RTK ligand genes induced by G12C-I treatment were similar, but not identical, to those evoked by MEK-I (Figure 3A). Analogous results were obtained by in studies of ARS-treated mouse KCP cells (Supplemental Figure 3A-B).

**Figure 3.**
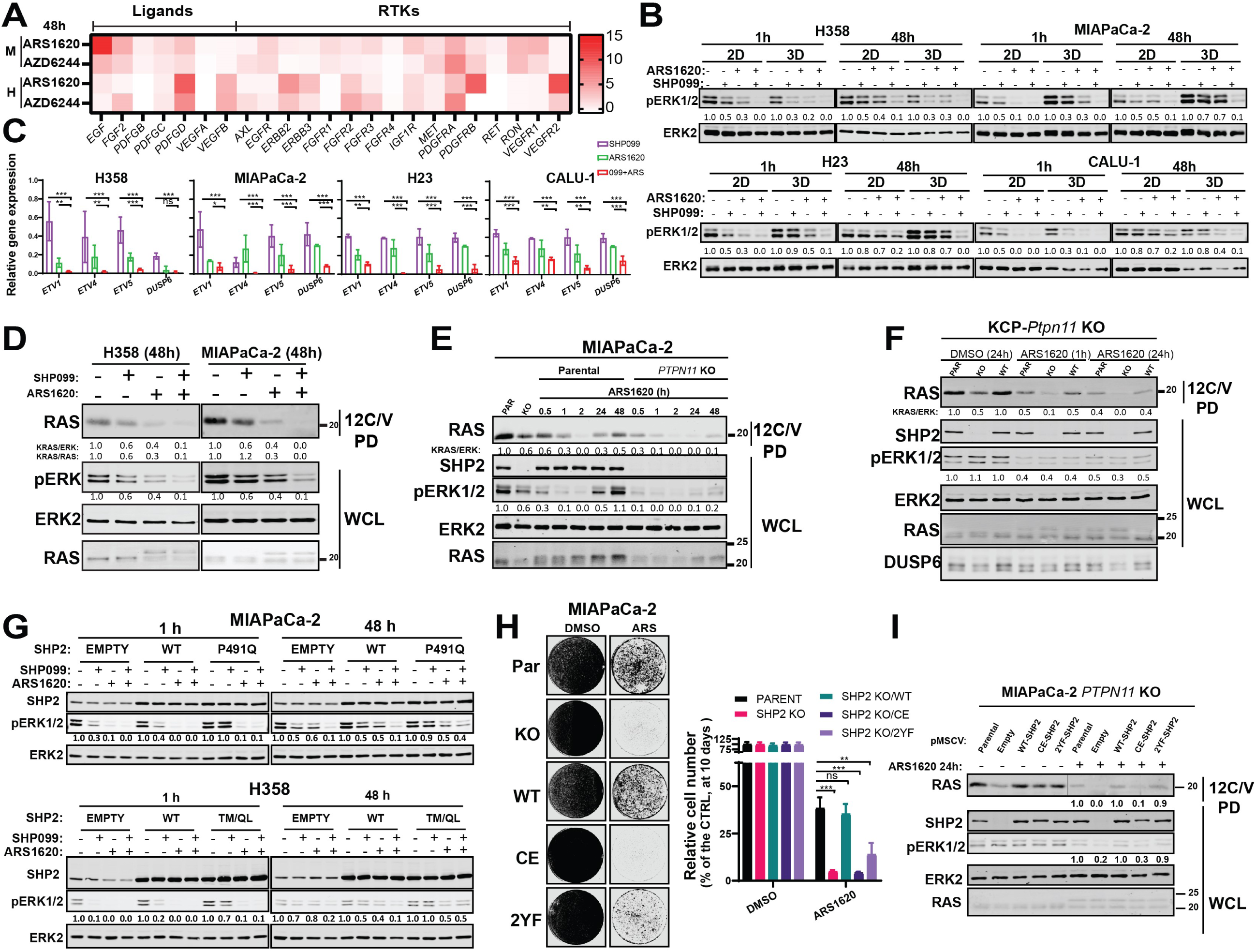
SHP2 inhibition acts upstream of RAS to abrogate G12C-I–evoked ERK MAPK pathway reactivation. **A**, Heat map showing time-dependent increase in RTK and RTK ligand gene expression in MIAPaCa-2 (M) and H358 (H) cells after DMSO, ARS1620, or AZD6244 treatment for 48h, as determined by qRT-PCR. **B**, Immunoblots of whole cell lysates from the indicated KRAS^G12C^-expressing cells treated with DMSO, SHP099, ARS1620, or both drugs (Combo) for 1h or 48h in 2D or 3D conditions. **C**, ERK-dependent gene expression (*ETV1, 4, 5* and *DUSP6*), assessed by qRT-PCR, in KRAS^G12C^ lines treated as indicated. **D**, SHP099 blocks RAS/ERK reactivation after 48h ARS1620 treatment of H358 and MIAPaCa-2 cells, assessed by 12C/V MB pulldown (PD) from whole cell lysates (WCL) **E**, Immunoblots of whole cell lysates (WCL) and 12C/V MB pull-downs (PD) from parental or *PTPN11*-KO MIAPaCa-2 cells treated with ARS1620 for the indicated times. **F**, Immunoblots of WCL and 12C/V MB pull-downs (PD) from parental KCP cells or *Ptpn11*-KO KCP cells with or without reconstituting with WT-*PTPN11*, treated with ARS1620 for the indicated times **G**, Immunoblots for SHP2, pERK, and ERK in WCL from MIAPaCa-2 and H358 cells ectopically expressing wild-type SHP2 (WT) or an SHP099-resistant mutant (P491Q or T253M/Q257L respectively), treated as indicated. **H**, Colony formation assay (12 days) on parental, *PTPN11* KO MIAPaCa-2 cells or *PTPN11* KO MIAPaCa-2 cells reconstituted with WT, phosphatase-inactive C459E (CE), or C-terminal tyrosine phosphorylation site-defective Y542F+Y580F (2YF) *PTPN11* mutants, with or without ARS1620 treatment. **I**, Immunoblots of WCL and 12C/V MB pull-downs (PD) from parental, *PTPN11* KO-MIAPaCa-2 or *PTPN11* KO reconstituted with WT, C459E (CE), Y542F+Y580F (2YF) *PTPN11* treated with ARS1620 for the indicated times. The images shown are representative of at least two independent biological replicates. For all experiments, drug doses were: SHP099 10 μm/l, ARS1620 10 μm/l, Combo = SHP099 10 μm/l + ARS1620 10 μm/l. Data represent mean ± SD; *P < 0.05; **P < 0.01; ***P < 0.001, two-tailed t test. Numbers under blots indicate relative intensities, compared to untreated controls, quantified by LICOR.

To probe the mechanism of adaptive resistance to G12C-Is, we assessed the biochemical effects of each single agent and the drug combination on RAS/ERK pathway activity after brief (1h)- and longer-term (48h) treatments. Single agent ARS blocked ERK1/2 phosphorylation after 1h in cells grown in 2D or 3D, but these effects were abolished after 48h of continued treatment (Figure 3B). Addition of fresh ARS to MIAPaCa-2 cells after 48h did not prevent pERK rebound (Supplemental Figure 3C), indicating that loss of pathway inhibition did not reflect drug metabolism or instability. By contrast, co-administration of SHP099 prevented the adaptive increase in ERK phosphorylation in response to ARS (Figure 3B and S3C). ERK-dependent gene expression often provides a better assessment of RAS/ERK pathway output than p-ERK levels (54), so we also measured key ERK-dependent genes in a panel of human G12C lines (by qRT-PCR) and in KCP cells (by RNAseq). Compared with either single agent, SHP099/ARS caused substantially greater suppression of ERK-dependent transcripts (Figure 3C and Supplemental Figure 3D).

Furthermore, 12C/V-MB PDs showed that mutant KRAS is reactivated upon 48h treatment with ARS; presumably, so are endogenous WT RAS isoforms. SHP099 blocked the adaptive increase in KRAS^G12C^-GTP (Figure 3D and Fig S3E), as did *SOS1* knockdown (Supplemental Figure 3F). *PTPN11* deletion had similar biochemical effects as SHP2 inhibition (Figure 3E), whereas re-expressing WT SHP2 restored adaptive resistance to ARS and sensitivity to SHP099 (Figure 3F and Supplemental Figure 3G), showing that the effects of SHP099 were on-target. The biochemical effects of SHP099 (like its effects on viability; Figure 1E-G) also were reversed in MIAPaCa-2 cells expressing *PTPN11*^*P491Q*^ (Figure 3G, Supplemental Figure 3H) and *PTPN11*^*T253M/Q257L*^-expressing H358 cells (Figure 3G). Hence, mutant KRAS is reactivated in G12C-I-treated cells, leading to RAS/ERK pathway re-activation, and SHP2-I, by acting upstream of SOS, blocks this adaptive response.

SHP2 catalytic activity is required for RAS/ERK pathway activation by most, if not all, RTKs, but its C-terminal tyrosyl residues are only essential in some RTK signaling pathways (55-57). Reconstituting *PTPN11*-knockout MIAPaCa-2 cells with WT *PTPN11*, but not a phosphatase-inactive mutant, *PTPN11*^*C459E*^(CE), restored ARS-induced adaptive resistance. SHP2 lacking both C-terminal tyrosine phosphorylation sites (*PTPN11*^*Y542F/Y580F*^, 2YF) partially restored adaptive resistance (Figure 3H-I and Fig S3 I). Thus, as in RTK signaling, PTP activity is essential, whereas C-terminal tyrosine residues play a modulatory role, in ARS-invoked activation of RTK signaling.

### Combined SHP2/ARS inhibition is efficacious in PDAC models *in vivo*

We next assessed the effects of ARS (200mg/kg/d), SHP099 (75mg/kg/d) or SHP099/ARS on orthotopic KCP tumors established in syngeneic C57BL6 mice (35). Tumors were allowed to grow for 14 days, 4 mice were sacrificed to obtain baseline tumor sizes (average 100 mm^3^), and the rest were treated with each single agent or SHP099/ARS for 3 or 10 days, respectively. After 10 days, control tumors (Veh) had quadrupled in mass compared with the pre-treatment baseline. Single agents had a largely static effect, although SHP099 treatment was more efficacious. By contrast, all tumors in the SHP099/ARS arm regressed markedly, and treated mice showed no evidence of toxicity (Figure 4A, Supplemental Figure 4A, and Table S2). Immunoblot analysis of tumor lysates after 3 days of treatment revealed greater inhibition of KRAS^G12C^, pERK, and the ERK-induced protein DUSP6 following combination treatment (Figure 4B). RNAseq showed that, compared with vehicle or either single agent, SHP2-I/G12C-I enhanced the suppression of ERK-, MYC-, anti-apoptotic-, and cell-cycle genes, while increasing pro-apoptotic gene expression (Figure 4C and Supplemental Figure 4B-D). Notably, single agent ARS inhibited pERK and the expression of ERK-dependent genes at least as well as did SHP099. These findings, along with the greater efficacy of SHP099 vs. ARS raised the possibility of additional effects of SHP2 inhibition either in tumor cells themselves or on the TME. RNAseq (on tumors after 3 days of treatment) also showed that several RTKs and RTK ligands were induced by G12C-I, demonstrating that adaptive resistance via RTK over-activation occurs *in vivo*. SHP099 also induced several RTK/RTK ligands, but although there was substantial overlap with ARS effects, several of these genes were affected differentially (qualitatively and quantitatively) by each agent (Figure 4D).

**Figure 4.**
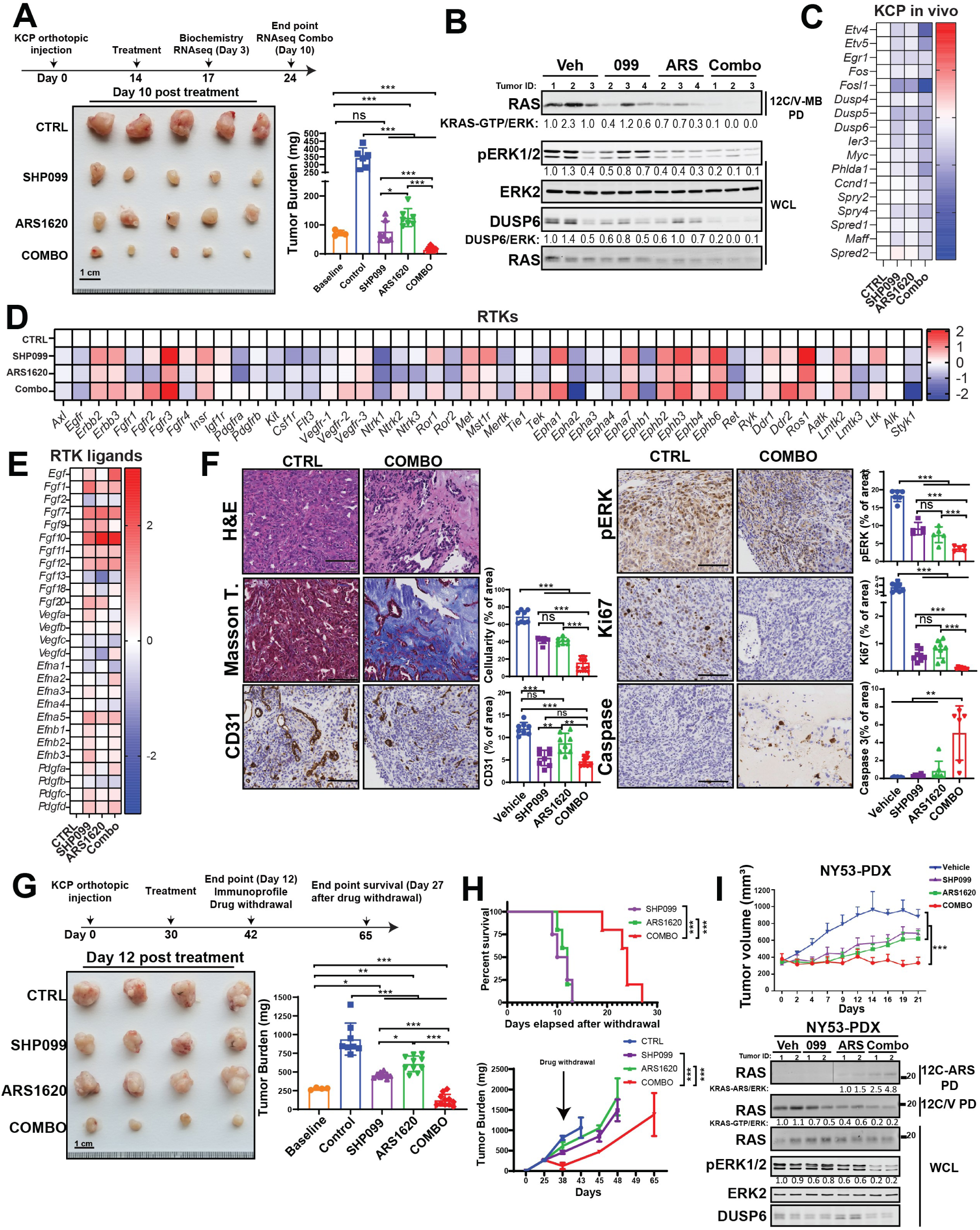
Combined ARS1620/SHP2 inhibition is highly efficacious in PDAC models *in vivo*. **A**, Pancreas tumors were established in syngeneic mice by orthotopic injections of KCP cells, and 14 days later, mice were treated with vehicle, SHP099, ARS1620 or both drugs (Combo), as depicted. Tumor weight was quantified in a cohort at Day 0 (baseline) and in treated mice at Day 10. **B**, Immunoblots of KCP-derived tumor lysates showing effects of the indicated treatments on KRAS^G12C^-GTP, pERK, and DUSP6 levels. **C**, ERK-dependent gene expression, assessed by RNAseq, in KCP tumors treated for 3 days, as indicated in **A** (colors indicate log2FC). **D-E**, Time-dependent increase in RTK (D) and RTK ligands (E) gene expression in KCP-derived orthotopic tumors after vehicle, SHP099, ARS1620 and Combo treatment at Day 3, determined by RNAseq (colors represent log2FC). **F**, H&E, Masson Trichome, CD31, pERK, Ki67 and cleaved Caspase 3 staining and quantification in KCP tumor sections from mice after 10 days of treatment, as indicated. **G**, KCP tumors were established in syngeneic mice and allowed to grow to much larger size before treatments were initiated, as depicted in the scheme. Tumor weight was quantified in one cohort before treatment, in another cohort after 12 days of treatment, and after drug withdrawal, at Day 27, as indicated. **H**, Kaplan-Meier curve of KCP tumor-bearing mice after withdrawal of the indicated drugs (top). Tumor growth curve after withdrawal of indicated treatment at day 12 (bottom). **H**, Response of sub-cutaneous NY53 patient-derived xenograft to treatment with vehicle, SHP099, ARS1620 or both drugs. For all experiments, drug doses were: SHP099 (75 mg/kg body weight, daily), ARS1620 (200 mg/kg body weight, daily) or both drugs (daily). Data represent mean ± SD; *P< 0.05, **P< 0.01, ***P < 0.001, one-way ANOVA with Tukey’s multiple comparison test. For K-M curves, log rank-test was used. Numbers under blots indicate relative intensities, compared with untreated controls, quantified by LICOR.

H&E-stained sections of tumors from SHP099/ARS-treated mice revealed a marked increase in collagenized stroma with scattered histiocytes, histiocytic giant cells, hemosiderin-laden histiocytes, and lymphocytes, compared with vehicle- or single agent-treated mice. Residual carcinoma cells were widely spaced with scattered glands in Combo-treated tumors, in contrast to the solid sheets of malignant cells seen in Control tumors. Masson-Trichrome staining confirmed the marked increase in collagen and diminished cellularity (Figure 4F and Supplemental Figure 4E). As we reported previously for KPC tumors (35), SHP099 treatment decreased KCP tumor vascularity, as shown by CD31 immunostaining (Figure 4F and Supplemental Figure 4D). Also similar to our previous findings on the effects of SHP099/MEK-I on KPC tumors, residual tumor cells in SHP099/ARS-treated (but not single agent- or vehicle-treated) mice showed evidence of ductal/acinar differentiation (Supplemental Figure 4E). Consistent with these observations, ductal and acinar, as well as endocrine, genes were induced (Supplemental Figure 4F). Immunohistochemical (IHC) analysis confirmed more profound pERK inhibition in tumors from SHP099/ARS-, compared with single agent-, treated mice (Figure 4F and Supplemental Figure 4D, along with decreased proliferation and increased apoptotic cell death (Figure 4F and Supplemental Figure 4D).

To evaluate efficacy even more stringently, we allowed KCP tumors to grow to 250mm^3^ before beginning treatment. Single agents again inhibited tumor growth, whereas SHP099/ARS caused dramatic tumor regression (Figure 4G). When treatment was stopped after 12 days, tumors recurred in all groups, but regrowth of SHP099/ARS-treated tumors was delayed, and median survival of this cohort more than doubled compared with single agent-treated mice (Figure 4H). SHP099/ARS treatment also was more effective than single agent treatment in inhibiting tumor growth and RAS/ERK pathway activation in a highly aggressive *KRAS*^*G12C*^ PDX model (Figure 4I).

### SHP099/ARS1620 evokes an anti-tumor immune program that is enhanced by anti-PD-1 therapy

The TME of human and mouse PDAC features abundant immune-suppressive myeloid cells, regulatory T cells (Tregs), and a scarcity of cytotoxic lymphocytes (58, 59). Consequently, these tumors are sometimes termed immunologically “cold.” Targeted therapies not only affect cancer cells, but most act on cells in the TME. SHP2 and the RAS/ERK pathway have roles in most, if not all, TME cells, often in several signaling pathways in these cells. G12C-Is, owing to their mutant specificity, only affect cancer cell signaling, but by altering growth factor/cytokine/chemokine production, they too could affect the TME of PDAC.

We used flow cytometry to survey the immune composition of the KCP TME, using the tumors from Figure 4G. Although the %CD45+ cells in tumors was unchanged in ARS-, SHP099-, and SHP099/ARS-treated mice, the composition of the CD45+ population was altered significantly. Total T lymphocytes (as % live cells) were increased in single agent- and, more substantially, in SHP099/ARS-treated groups (Figure 5B and Supplemental Figure 5A). Each single agent also increased B lymphocytes, and this increase was preserved in SHP099/ARS-treated mice (Figure 5B). By contrast, there was a trend towards decreased total CD11b+ myeloid cells, mostly comprising g-MSDC, following SHP099 alone (P=0.11), and a nearly 50% decrease (P=0.045) after SHP099/ARS (Figure 5B and Supplemental Figure 5B). Subset analyses revealed that single agent, and especially combination, treatment preferentially increased CD8 T cells (as % total T cells), but these cells exhibited markers (TIM3, PD1, OX40) consistent with “exhaustion” (Figure 5C). By contrast, CD4 cells and Tregs decreased, and consequently, CD8/Treg ratios increased, most prominently in response to SHP099/ARS (Figure 5D-E). We also examined the spatial distribution of immune cells in the TME by multi-color immunofluorescence (IF) and IHC in four tumors. The results comported with the flow cytometric data, although there were regional differences in lymphoid infiltration into tumors. Notably, SHP099/ARS-treated samples clearly had more intratumor lymphocytes (Supplemental Figure 5C).

**Figure 5.**
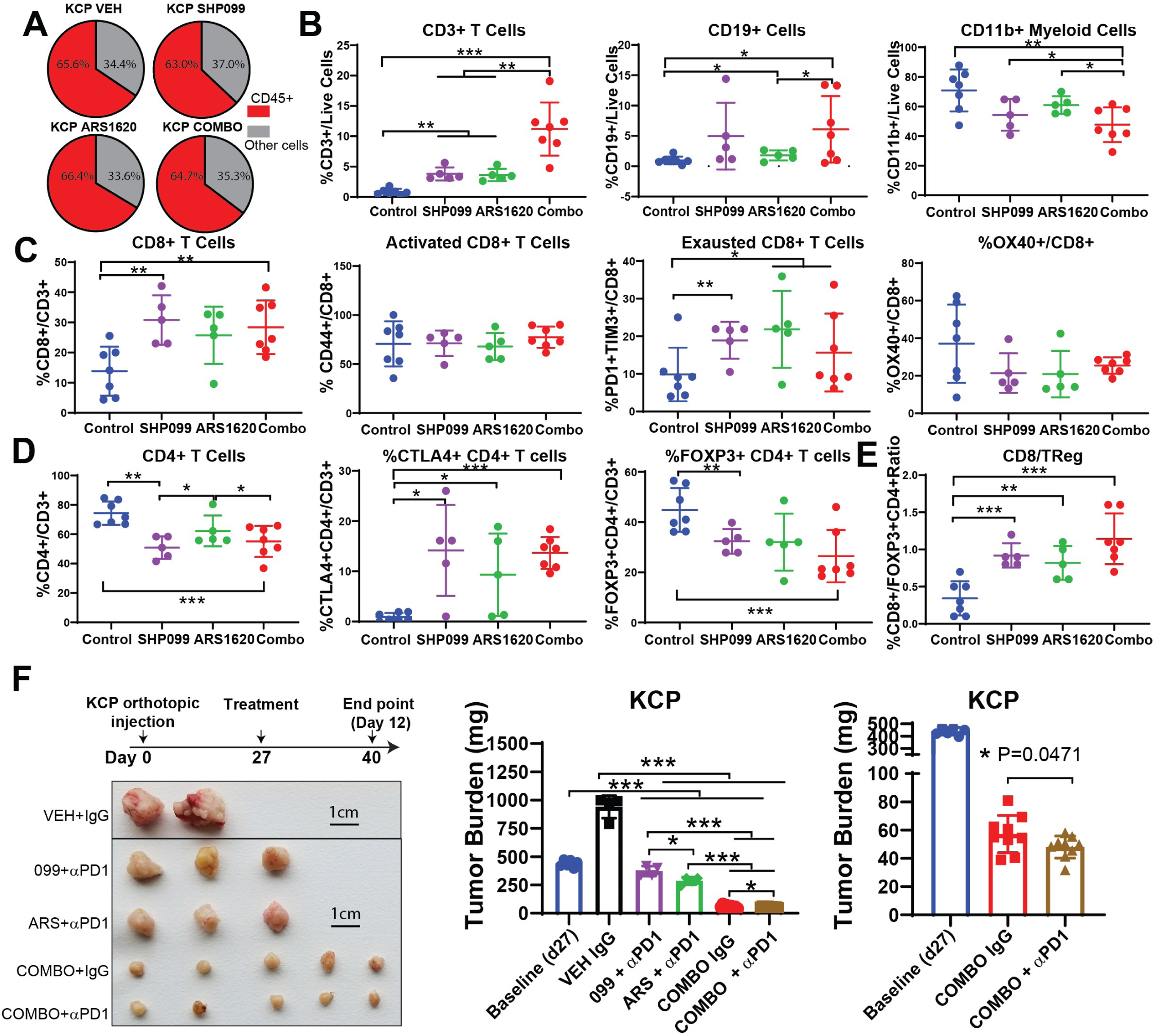
ARS1620/SHP099 combination provokes an anti-tumor immune program in syngeneic PDAC model that is enhanced by anti-PD-1. **A**, Pie charts showing %CD45+ (immune) cells and %CD45- (cancer plus stromal) cells in KCP-derived tumors treated for 12 days, as in Figure 4. **B**, Frequencies of infiltrating CD3+ T cells, CD19+ B cells, and CD11b+ myeloid cells **C**, Frequencies of infiltrating CD8+ T cells and respective sub-populations **D**, Frequencies of infiltrating CD4+ T cells and respective subpopulations **E**, Ratio of infiltrating CD8+ T cells to FOXP3+ regulatory CD4+ T cells. Data represent mean ± SD; \**P* < 0.05, \*\**P* < 0.01, \*\*\**P* < 0.001, multiple unpaired Welch’s *t* test (two tailed) For **B-E**, Tumor-infiltrating immune cells were analyzed at day 12 after the indicated treatments. **F**, Syngeneic mice bearing KCP tumors were treated with vehicle + isotype IgG (200 μg/mouse three times/week), SHP099 (75 mg per kg body weight, daily) + anti PD1 (200 μg/mouse three times/week), ARS1620 (200 mg per kg body weight, daily) + anti PD1 (200 μg/mouse three times/week), ARS1620+SHP099 (COMBO) (daily) + isotype IgG (200 μg/mouse three times/week) or COMBO (daily)+ anti-PD1 (200 μg/mouse three times/week), as depicted in the scheme. Tumor weight was measured at Day 0 (baseline) and Day 12. Right-most panel shows expanded scale for the indicated treatments. Data represent mean ± SD; \**P* < 0.05, \*\**P* < 0.01, \*\*\**P* < 0.001; one-way ANOVA with Tukey’s multiple comparison test.

Single agent AMG510 caused an ∼50 fold increase in intra-tumor CD3+ and CD8+ T cells in sub-cutaneous (SQ) xenografts of *KRAS*^*G12C*^-engineered CT26 colon cancer cells (20). By contrast, we observed only a modest increase in T cells in ARS-treated orthotopic KCP tumors (Figure 5 B-C). To ask whether this difference might reflect the distinct location of the tumors in each study, we compared the efficacy of SHP099/ARS in mice with orthotopic (ORTHO) or SQ KCP tumors. Indeed, SQ tumors showed greater regression and exhibited a more robust anti-tumor T cell immune-response (Supplemental Figure 5D-E).

SHP099 and G12C-I each increased the expression of chemokine and cytokine genes that promote T cell recruitment (e.g., *CXCL9-11; CCL5*) (60-64), while decreasing the expression of those (e.g., *CXCL1-5; CCL9*) (61, 63, 65, 66) that favor immune-suppressive CD11b+ myeloid cells (Supplemental Figure 6C). SHP099/ARS evoked much greater differential expression of these genes, which likely accounts, at least in part, for the more favorable immune modulatory effects of the combination. The increase in PD1+ (potentially “exhausted”) T cells in SHP099/ARS-treated mice suggested a possible benefit of adding PD-1 blockade to this regimen. Indeed, SHP099/ARS/PD1 resulted in even greater tumor regression compared with SHP099/ARS or either single agent combined with anti-PD1 (Figure 5F). H&E- and Masson Trichrome-stained sections of tumors from SHP099/ARS/PD1-treated mice revealed large areas of collagen scarring and only scattered residual tumor cells (Supplemental Figure 6 A-B). Thus, SHP099/ARS increases T cell infiltration and reactivity in previously immunologically “cold” tumors and sensitizes PDAC to immune checkpoint blockade.

### Effects of SHP2 inhibition in PDAC cancer cells and tumor microenvironment

To begin to assess which of the above findings reflected direct effects of SHP099 on tumor cells (vs. indirect effects of SHP2 inhibition on TME cells), we established orthotopic tumors of *Ptpn11-*KO KCP cells reconstituted with *PTPN11* or the drug resistant mutant *PTPN11*^*T253M/Q257L*^ (TM/QL). SHP099 treatment (for 10 days) suppressed the growth of *PTPN11*-reconstituted KCP tumors, but TM/QL-reconstituted tumors showed no evidence of regression (Figure 6A). As in parental KCP tumors, SHP099 evoked an influx of CD8 and CD4+ T cells in *PTPN11*-reconstituted KCP tumors. This influx also was abrogated in TM/QL-KCP tumors (Figure 6B-C). RNAseq revealed a failure of TM/QL tumors to alter the expression of the chemokines and cytokines that likely mediate observed T cell infiltration in mice bearing parental KCP tumors or WT-reconstituted, KCP-KO tumors (Supplemental Figure 6C). These data demonstrate that SHP099 (and, presumably, SHP099/ARS) must alters signaling and induces cell death in KCP tumors to evoke change in the immune microenvironment (although additional effects of SHP099 on immigrating immune cells might also be needed for the anti-tumor response; see Discussion).

**Figure 6.**
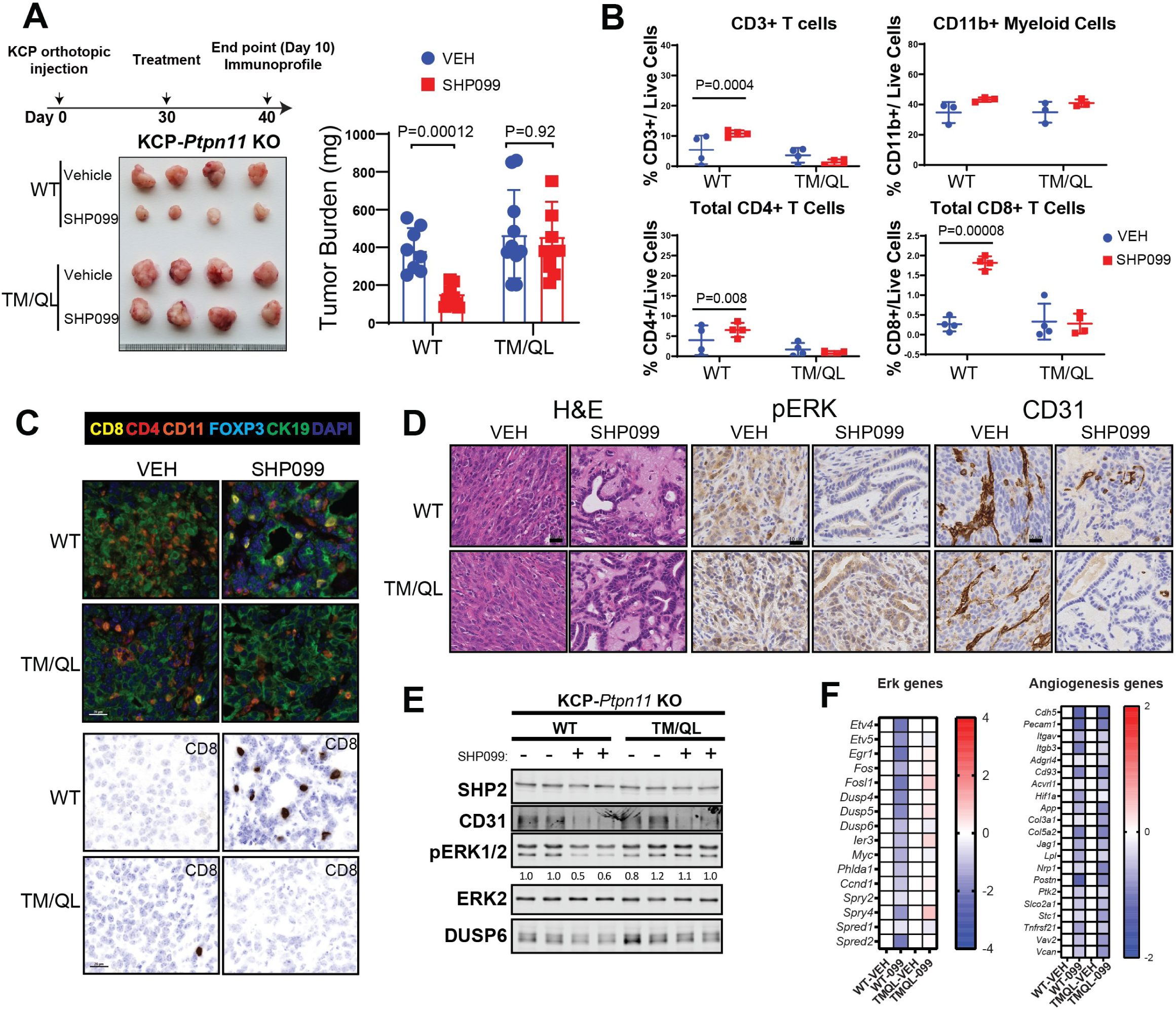
Tumor cell-autonomous and non-autonomous effects of SHP2 inhibition in PDAC. **A**, Tumors were established in syngeneic mice by orthotopic injections of *Ptpn11* KO-KCP cells reconstituted with wild-type (WT) or SHP099-resistant *PTPN11*^T253M/Q257L^ mutant (TM/QL) and treated with vehicle or SHP099 (75 mg/kg body weight, daily), as depicted in the scheme. Tumor weights were measured at Day 10. **B**, Tumor-infiltrating immune cells from experiment in A (n=4). **C**, Multiplex IF/IHC analysis of representative tumors from A. **D**, H&E, pERK and CD31 staining from representative KCP tumors, as described in A (n=6). **E**, Immunoblot showing CD31, pERK, and DUSP6 levels in representative tumors from A. **F**, ERK- and angiogenesis-dependent gene expression, assessed by RNAseq, in KCP tumors from experiment described in A (n=5) (colors indicate log2FC). Data represent mean ± SD; Significance was assessed by multiple unpaired Student’s *t* test (two-tailed).

Although SHP099 treatment did not decrease the size of TM/QL tumors, histological examination surprisingly showed areas of tumor necrosis, replacement with eosinophilic material, and more duct-like epithelial architecture (Figure 6D). IHC revealed clear rescue of pERK staining, while RNAseq showed that ERK target gene expression was unaffected in TM/QL tumors from SHP099-treated mice, confirming that this mutant was, as expected, SHP099-resistant and that TM/QL tumor cells were unaffected by SHP099. Nevertheless, TM/QL-KCP tumors, like their WT-reconstituted (and parental KCP) counterparts, showed marked hypo-vascularity, which was confirmed by reduced CD31 staining and decreased expression of angiogenic genes (Figure 6D-F and Fig S6D-E). Therefore, in addition to its effects on tumor cells, SHP099 has direct anti-angiogenic actions. SHP099 treatment also reduced the number of activated fibroblasts in KCP-WT tumors, as confirmed by αSMA staining and *Acta2* expression. However, these effects were reversed in SHP099-treated TM/QL-KCP-tumor-bearing mice (Supplemental Figure 6F-G). Hence, the ability of SHP099 to modulate tumor-associated fibrosis requires inhibitor action in tumor cells, presumably to decrease the production of secreted factors that act on stromal fibroblasts. Intriguingly, *Fgf2* expression was decreased after SHP099 treatment in WT-KCP-tumor-bearing mice., while this effect was rescued in treated TM/QL-KCP-tumor-bearing mice (Supplemental Figure 6G), suggesting that this growth factor might be particularly important for tumor-associated fibrosis in this model.

### ARS1620/SHP099 is also efficacious in *KRAS*^*G12C*^ NSCLC

To ask if the efficacy of SHP099/ARS for targeting *KRAS*^*G12C*^-mutant tumors extended to other histotypes, we monitored the effects of SHP099, ARS, and SHP099/ARS on *KRAS*^*G12C*^ (KC) and *Kras*^*G12C*^; *Tp53*^*R270H*^ (KCP) NSCLC GEMMs (67) by serial magnetic resonance imaging (MRI). SHP099/ARS induced deep responses by 2 weeks, and complete responses (CRs) after 4 weeks, of treatment. Remarkably, all combination-treated mice remained in remission over an 8-week treatment period (Figure 6A-B). Mice bearing KC or KCP NSCLC and treated with either single agent had much shallower initial responses. Whereas SHP099-treated KC-tumors remained in remission, tumors recurred in single agent ARS-treated mice by 8 weeks on therapy, and KCP tumors recurred by 8 weeks of treatment with either single agent. Consequently, while ARS or SHP099 treatment resulted in a marginal survival advantage, 100% of SHP099/ARS-treated mice remained alive and disease-free throughout the treatment period (Figure 7E-F). SHP099/ARS also inhibited the growth of *KRAS*^*G12C*^-driven H2122 xenografts more effectively than either single agent (Supplemental Figure 7A).

**Figure 7.**
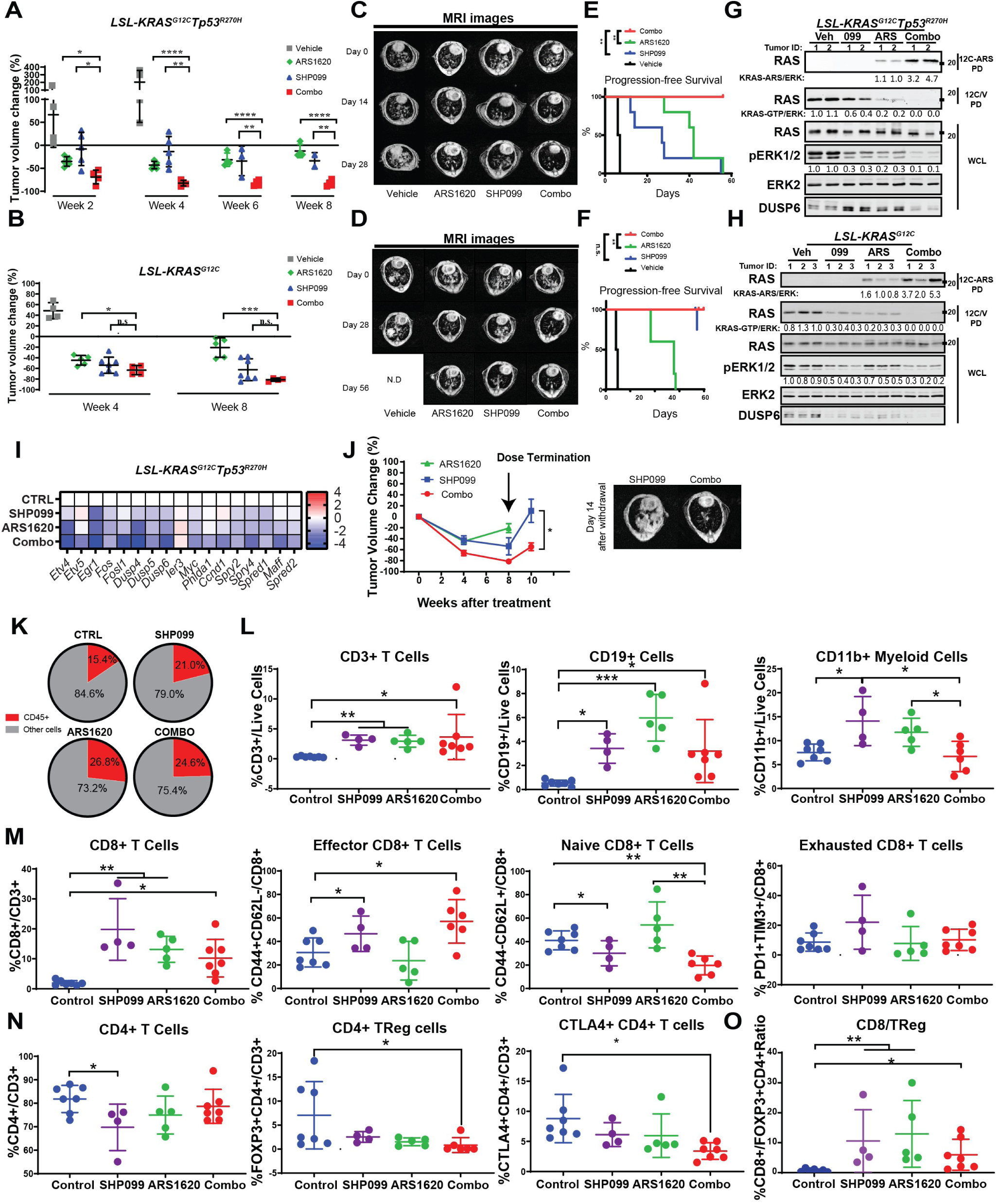
ARS1620/SHP099 combination is also efficacious in NSCLC GEMM. **A-B**, Change in tumor volume in *LSL-KRAS*^*G12*C^-*Trp53*^*R270H*^ (A) and *LSL*-*KRAS*^*G12C*^ (B) NSCLC GEMMs, quantified by MRI, after treatment with vehicle, SHP099, ARS1620 or both drugs (Combo) at the indicated times; Data represent mean ± SD; \**P* < 0.05, \*\**P* < 0.01, \*\*\**P* < 0.001, \*\*\*\**P* < 0.0001; one-way ANOVA with Tukey’s multiple comparison test. **C-D**, Representative MR images show lungs from *LSL-KRAS*^*G12C*^*-Trp53*^*R270H*^ (C) and *LSL-KRAS*^*G12C*^ (D) NSCLC GEMMs before and after treatment, as indicated. **E-F**, Kaplan-Meier curves for *LSL-KRAS*^*G12C*^*-Trp53*^*R270H*^ (E) and *LSL-KRAS*^*G12C*^ (F) models after the indicated treatments; \**P* < 0.05, \*\**P* < 0.01, \*\*\**P* < 0.001 log-rank test. **G-H**, Immunoblots of lysates and 12C/V-MB or 12C-ARS Fab pull-downs (PD) from *LSL-KRAS*^*G12C*^*-Trp53*^*R270H*^ (G) and *LSL-KRAS*^*G12C*^ (**H**) tumors after 3 days of treatment. **I**, ERK-dependent gene expression, assessed by RNAseq, in tumors from *LSL-KRAS*^*G12C*^*-Trp53*^*R270H*^ mice, treated for 3 days, as indicated (colors indicate log2FC). **J**, *LSL-KRAS*^*G12C*^ tumor volume after treatment and drug withdrawal, as indicated (left); representative MR images 14 days after drug withdrawal are shown at right. **K**, Pie chart showing %CD45+ and %CD45-cells in *LSL-KRAS*^*G12*C^ tumors after 6 days of treatment, as indicated (data are presented as the average of each treatment, n=5 or 7) **L**, Frequencies of infiltrating CD3+ T cells, CD19+ B cells, and CD11b+ myeloid cells **M**, Frequencies of infiltrating CD8+ T cells and respective sub-populations **N**, Frequencies of infiltrating CD4+ T cells and respective subpopulations **O**, Ratio of infiltrating CD8+ T cells to FOXP3+ regulatory CD4+ T cells. For **L-O**, The indicated cell populations from *LSL-KRAS*^*G12C*^ tumors were analyzed after 6 days of treatment. Data represent mean ± SD; \**P* < 0.05, \*\**P* < 0.01, \*\*\**P* < 0.001, multiple unpaired Welch’s *t* test (two-tailed).

Biochemical analysis revealed similar G12C inhibition by SHP099 or ARS, but substantially greater inhibition with SHP099/ARS. Direct measurements, using the 12C-ARS-Ab, confirmed increased adduct formation in tumors from SHP099/ARS-, compared with single agent ARS-treated mice. As in the PDAC model, RNAseq (at day 3) in KCP-treated mice, showed induction of several RTKs and RTK ligands by G12C-I treatment, confirming that adaptive resistance via upregulation of RTK signaling also occurs in NSCLC (Supplemental Figure 7D-E). Addition of SHP2 inhibitor abrogated this resistance, as shown by the greater suppression of pERK and ERK-, MYC-, apoptotic-and cell cycle-target gene expression evoked in SHP099/ARS-compared with ARS- (or SHP099-) treated mice (Figure 7I and Supplemental Figure 7B-F).

When treatment of KC tumor-bearing mice was stopped at 8 weeks, tumors recurred in SHP099- and SHP099/ARS-treated mice. Again, however, recurrence was substantially slower in the latter group, indicative of substantially fewer surviving tumor cells (Figure 7J). We also tested the effects of the clinical grade G12C-I, MRTX1257 (MRTX), alone or combined with SHP099, on KC tumors. Although, consistent with its higher potency, single agent MRTX was more efficacious than ARS in this model, SHP2 inhibition also enhanced MRTX efficacy, as revealed by the slower tumor recurrence after drug withdrawal (Supplemental Figure 7G).

We noticed that, as in PDAC, disease control was better in SHP099-treated than in ARS-treated KC tumor-bearing mice, despite a comparable decrease in KRAS^G12C^-GTP and, if anything, greater suppression of ERK target genes in ARS-treated mice (Fig 7B and D). These data again suggested additional effects of SHP2-inhibition, potentially on the TME. To investigate the phenotypic and functional alterations of immune cells, we harvested mouse tumor-bearing lungs after 6 days of treatment. Flow cytometry revealed enhanced immune cell infiltration into tumors from ARS-, SHP099-, and SHP099/ARS-treated KC mice (Figs. 7K and S8A). Similar to the effects on PDAC, we found that total T and B lymphocytes (as % live cells) were increased in single agent- and SHP099/ARS-treated groups (Figure 7L). Subset analyses of the CD3+ population revealed a relative increase in CD8+ T cells for in all treatment groups. Profiling of CD8+ T cells showed a substantial increase of CD44+CD62L-effector CD8+ T cells after SHP099 and SHP099/ARS treatment, consistent with a respective decrease in naïve cells (CD44-CD62L+) (Figure 7M). However, in contrast to its effects in PDAC, ARS-, SHP099-, and SHP099/ARS revealed no change in PD1+ TIM3+ CD8+ cells in any group (Figure 7M). Single agent SHP099 led to decreased tumor infiltration of CD4+ T cells, but this decrease was mitigated by ARS addition. Profiling of CD4+ T cells showed that SHP099/ARS decreased Treg cells (FOXP3+), resulting in increased CD8/Treg ratio (Figure 7N-O). Enhanced T cell infiltration into KC and KCP tumors from SHP099/ARS-treated mice was also evident by multiplex IF/IHC (Supplemental Figure 8B).

Concomitant with these potentially beneficial effects on tumor-associated T cells, there were complex effects on tumor myeloid cells. Each single agent increased total CD11b+ cells, and the fraction of this compartment composed of macrophages (F4/80+Gr1-) (Figure 7L and Supplemental Figure 8B). SHP099/ARS reversed this increase and led to slight overall decrease in total CD11b+ cells and macrophages (Figure 7L and Supplemental Figure 8B). There was no change in m-MDSCs (CD11b+Ly6C+LY6G-), although there was increase in total g-MDSCs (CD11b+Ly6C-LY6G+) (as % live cells) (FigureS8C). Dendritic cells (CD11c+F4/80-GR1-) were also significantly reduced after SHP099/ARS. Taken together, these data show that SHP099/ARS combinations also have beneficial effects in NSCLC model and suggest additional rational combinations for enhancing their efficacy.

Finally, we assessed the effects of SHP099, ARS, and SHP099/ARS on tumor-associated vasculature. In contrast to the effects of SHP099 in PDAC (Figure 4F and Supplemental Figure 4D), in the KC and KCP NSCLC models, both single agents and SHP099/ARS increased lung tumor-associated blood vessels (Supplemental Figure 7H-I). These data indicate that SHP099 (and, presumably, SHP099/ARS) might trigger a tumor cell autonomous secretory program that promotes tumor angiogenesis specifically in the NSCLC TME (see Discussion).

## DISCUSSION

The advent of clinically active, covalent KRAS^G12C^ inhibitors provided the first opportunity to directly target this key driver oncoprotein in tumors (20, 22, 68). However, initial reports from phase I trials show tumor responses in *KRAS*^*G12C*^ tumors are partial and restricted to a subset of patients (20, 22). These results suggest that, similar to other “targeted therapies” (51, 69, 70), G12C-Is, as single agents, will have limited impact due to drug resistance. “Adaptive resistance,” in which the inhibited pathway is reactivated due to induction of genes for RTKs/RTK ligands, is a common form of intrinsic resistance to targeted agents (48, 49, 71-75). The host immune system can cure some malignancies; conceivably, all cancer cures might require generation of a durable anti-tumor response. However, most tumors evade anti-tumor immunity via diverse mechanisms. Furthermore, conventional chemotherapy and most targeted therapies also affect the TME, and these actions are not often considered carefully. A sophisticated approach to developing curative cancer regimens requires delineating the mechanism of action of anti-neoplastic agents on cancer and TME cells and using these insights to develop complementary combinations that prevent tumor resistance. Here, by using two new affinity reagents that allow direct monitoring of KRAS^G12C^ activation and inhibition, we find that G12C-Is evoke adaptive resistance *in vitro* and *in vivo* by inducing KRAS^G12C^ re-activation. Similar to the effects of other RAS/ERK pathway inhibitors (48, 49, 71-75), G12C-Is induce RTK/RTK ligand genes, increasing RTK signaling to RAS. SHP2 inhibition, by increasing G12C-I accessibility to mutant KRAS, abrogates adaptive resistance. By studying *KRAS*^*G12C*^ PDAC and NSCLC GEMMs, we find that SHP2-Is and G12C-I/SHP2-I combinations have complex effects on the TME, some mediated indirectly via effects on tumor cells, but others that reflect direct effects on tumor endothelial cells. Our results suggest additional combination approaches to further enhance G12C-I efficacy.

While this work was in progress, others reported that G12C-Is induce adaptive resistance in G12C-mutant cell lines and PDXs. Although our findings agree generally with these studies, they differ in important details. Misale *et al*. (39) reported that ARS evokes adaptive resistance by activating the PI3K-AKT pathway. However, we did not observe increased activation of AKT (pAKT) or downstream targets (e.g., pS6) in ARS-treated H358 and MIAPaCa-2 cells or in KCP pancreas tumors from ARS-treated mice (data not shown). Ryan *et. al*. (40) argued that adaptive resistance to G12C-Is involves upregulation of RTK signaling and activation of WT RAS, which cannot be targeted by the inhibitor, whereas Xue *et al*. (41) reported that resistance arises from pre-existing heterogeneity that enables some tumor cells to survive by inducing mutant *KRAS* to levels that exceed inhibitor targeting capacity. Like Ryan *et al*., we observed induction of RTKs/RTK ligand genes following ARS treatment, and similar to our previous observations on MEK-I effects (35), we saw different patterns of RTK/RTK ligands induced by G12C-I in different cell lines, even within the same cancer histotype. We also noted qualitative and quantitative differences in RTK/RTK ligand gene induction by MEK-Is and G12C-Is (Fig 3A and data not shown); the mechanism underlying these differences merits future study. In contrast to the previous reports, however, we did not observe altered *KRAS* gene expression *in vitro* or *in vivo* in response to G12C-I treatment. Instead, by capitalizing on our novel 12C/V-MB PD assay, which enables specific monitoring of G12C-GTP (in the presence of normal RAS-GTP) *in vitro* and *in vivo*, we show clearly that adaptive resistance to ARS is accompanied by reactivation of KRAS^G12C^. Although normal KRAS/other RAS isoforms might also be reactivated, KRAS^G12C^-GTP (given its GAP resistance) should accumulate to higher levels than other RAS-GTP species and thus is likely to be the main effector of ERK MAPK pathway reactivation. Consistent with this analysis, we found that ARS induces adaptive resistance equivalently in *Kras*^*wt*^/*KRAS*^*G12C*^ and *Kras*^*-/-*^/*KRAS*^*G12C*^ MEFs (Supplemental Figure 1A). Our new affinity-capture approaches will enable similar assessment of G12C activation state in other systems.

Several lines of evidence show that SHP099 was “on-target” in our experiments, and hence that SHP099 action reflects SHP2 inhibition. First, SHP099 had the expected biochemical effects on the RAS/ERK pathway in the multiple lines tested. Moreover, two different drug-resistant mutants (*PTPN11*^P491Q^, *PTPN11*^*T253M/Q257L*^) rescued these effects (in PDAC *and* NSCLC cells, respectively). Furthermore, *PTPN11* KO or *SOS1* knockdown had biological and biochemical consequences similar to those of SHP099. Our results are consistent with, and strengthen, previous studies of the effects of SHP2 modulation on G12C-I inhibitor action (22, 38-41). *PTPN11*-KO cell reconstitution experiments (Figure 3H-I and Fig S3 I-J) revealed that PTP activity is essential, whereas C-terminal tyrosine residues play a modulatory role, in adaptive resistance to G12C-Is. These results comport with previous studies on SHP2 action in RTK signaling (55-57), which found that C-terminal phosphorylation is essential some, but not RTK pathways, and thus provide clues into which re-activated RTKs are most critical to mediating adaptive resistance.

We also find that SHP2 and/or G12C inhibition in immune-competent murine PDAC and NSCLC models has important effects both on tumor cells and cells in the TME. Similar to the initial reports of G12C-I and SHP2-I actions in the clinic (20, 22, 33), single agent ARS or SHP099 had limited efficacy in either tumor type (Figure 4A,G-H and Figure 7A-F). By contrast, G12C-I/SHP2-I dramatically improved efficacy and extended survival in all models tested without evident toxicity (Figure 4A, G-H and Figure 7A-F). Moreover, SHP099 abrogated adaptive resistance to ARS in tumors via the same mechanism observed *in vitro*: it increased occupancy of the KRAS^G12C^-GDP state facilitating greater ARS engagement, and thereby restored KRAS/ERK pathway inhibition, enhanced suppression of ERK-, MYC-, anti-apoptotic-, and cell-cycle genes, and concomitantly induced pro-apoptotic genes.

Although each single agent affected the TME, SHP2-I/G12C-I evoked a much broader immune response in both PDAC and NSCLC, featuring increased infiltration of CD8+T cells, decreased Tregs and consequent higher CD8/Treg ratio, and an increase in tumor-associated B cells. Nevertheless, the responses of the PDAC and NSCLC models diverged in important ways. For example, presumed immune-suppressive populations differed: CTLA4+/CD4+ T cells increased after SHP099, ARS, or SHP099/ARS treatment in PDAC (Figure 5D) but decreased in NSCLC (Figure 7N). Conversely, immune-suppressive CD11b+ myeloid sub-populations (m-MDSC, g-MDSC) generally decreased in PDAC (Figure 5A; Supplemental Figure 5B) but increased in NSCLC (Figure 7L; Supplemental Figure 8B). Furthermore, SHP2-I/G12C-I-induced CD8^+^ T cells displayed exhaustion markers only in the PDAC model (Figure 5C). Accordingly, in the PDAC model adding anti-PD-1 to ARS alone or SHP099/ARS enhanced anti-tumor immunity and conferred additional therapeutic benefit (Figure 5 F; Supplemental Figure 6A-B).

Gene expression analysis suggests key chemokines that likely mediate SHP2-I and G12C-I effects on the TME. Increased CXCL9-11 caused by SHP099, ARS, or SHP099/ARS treatment probably promote enhanced T cell immigration into orthotopic KCP tumors, whereas increased CXCL13 could evoke B cell immigration. By contrast, decreased *CXCL1-3/5* and/or *CCL9* could account for the altered myeloid cell population (Fig S5F-G). Notably, previous studies reported that activated KRAS promotes enhanced secretion of CCL9 and CXCL3, which recruit immunosuppressive macrophages and MDSCs. These actions were attributed to MYC activation or repression of IFN-regulatory factor (IRF)-2, respectively (65, 66). In line with these studies, SHP099, ARS, or SHP099/ARS decreased *Myc* levels (Figure 4C), while upregulating expression of *Irf1, 2, 7 and 9* (data not shown). Enhanced IRF activity might also explain the observed increase in *CXCL9-11* and *CXCL13* (76-78).

Recently, it was reported that AMG510 evoked an ∼50 fold increase in intratumor CD3+ and CD8+ T cells in SQ xenografts of *Kras*^*G12C*^-engineered CT26 CRC cells in syngeneic Balb/c mice (20). We observed much more modest increases in intratumor T cells in ARS-treated mice bearing orthotopic pancreas or autochthonous lung tumors. This discrepancy can be explained, at least in part, by differences in tumor location: SQ KCP tumors showed more robust T cell infiltration and greater anti-tumor responses than orthotopic tumors (Supplemental Figure 5D-E), in line with similar tumor site-dependent, quantitative and qualitative differences in response to other therapeutic modalities (59). We do not exclude the possibility that the precise genetics or histotype of CT26 and KCP tumors, the strain in which the tumors were established (Balb/c vs C57BL/6), and/or the respective potency of ARS vs AMG could also contribute. Regardless, these results emphasize the need to carefully investigate all of these parameters in credentialing single agents or combinations.

Perhaps surprisingly, single SHP099 was more efficacious than ARS alone both in PDAC and NSCLC, even though ARS directly targets the driver oncogene. ARS1620 is less potent than other G12C-Is, such as AMG-510 and MRTX1275 (20, 22). Nevertheless, we think that inadequate potency is unlikely to explain the superior single agent efficacy of SHP099 (compared with ARS) or the improved efficacy of SHP099/ARS combinations *in vitro* or *in vivo*. ARS was at least as effective as SHP099 in lowering KRAS^G12C^-GTP, pERK levels, and ERK-dependent gene expression *in vitro* (Figure 3D) and *in vivo* (Figure 4B, I; Figure7G and Supplemental Figure 7A). SHP099/ARS further enhanced suppression all of these parameters. SHP099 also potentiated the effects of AMG510 *in vitro* (Supplemental Figure 1K) and of MRTX1275 *in vivo* (Supplemental Figure 7G). Although we confirmed that AMG-510 and MRTX1275 had greater single efficacy than single agent ARS or SHP099 *in vitro* and *in vivo*, our results suggest that the superior anti-tumor effects of SHP099 compared with ARS are the result of SHP2 actions in cells in the TME.

By reconstituting *Ptpn11-*KO KCP cells with the drug-resistant mutant *PTPN11*^*T253M/Q257L*^, we could distinguish direct effects of SHP2 inhibition in tumor cells and cells in the TME, respectively. Indeed, SHP099-resistant KCP pancreas tumors showed marked decreases in tumor vasculature in response to SHP099 treatment, similar to their parental (35) and WT-*PTPN11*-reconstituted (Figure 6D)counterparts. This decrease in vascularity apparently reduced tumor perfusion, given the histological evidence of central necrosis and decreased tumor cell density (Figure 6D). Intriguingly, histology and gene expression analysis also suggest that TM/QL-tumors partially differentiated in SHP099-treated mice, even though the tumor cells themselves were unresponsive to SHP099 as confirmed by retained p-ERK levels and ERK-dependent gene expression. Whether these effects are due to altered production of a factor(s) from endothelial cells (or some other, as yet undefined, SHP099-sensitive component of the TME) or decreased autocrine signaling by the reduced tumor cell population remains unclear. Moreover, SHP099 effects on tumor vasculature are context-dependent, as it evokes a moderate increase of tumor vascularity in NCSCLC models (Fig S7 H-I). Such differences might arise and intrinsic endothelial cell heterogeneity (79, 80) and/or marked differences in oxygenation of PDAC and NSCLC.

By contrast, the SHP099-evoked influx of T cells and the decrease in activated fibroblasts seen in WT-*PTPN11* expressing tumors was abrogated in TM/QL-KCP tumors, indicating that SHP2 must be inhibited within tumor cells to evoke these changes. ARS only affects mutant KRAS, so its action on cells in the TME (and by inference, any additional effects it contributes to SHP099/ARS) must reflect altered mediator production by tumor cells, a notion supported by our RNAseq results (Supplemental Figure 6C). However, our findings do not exclude additional direct effects of SHP2 inhibition on signaling pathways in TME cells. Mice that express drug-resistant SHP2 in specific cells in the TME are required to address such issues; such studies are underway in our laboratory.

In summary, we find that G12C-I/SHP2-I efficacy derives from a combination of actions on tumor cells and cells in the TME, reveal direct anti-angiogenic effects of SHP2-Is, and demonstrate that G12C-I/SHP2-I can combine with PD1 checkpoint blockade to improve therapeutic outcomes in *KRAS*^*G12C*^ mutant tumors. In the NSCLC model, mice remained in remission for up to 8 weeks of continuous combination therapy. Nevertheless, after treatment cessation, both types of tumor relapsed. Our results also suggest additional rational combinations that might enhance efficacy and effect cure (e.g., SHP2-I/ G12C-I +anti CTLA4 in PDAC; SHP2-I/ G12C-I +g-MDSC-targeted therapy in NSCLC). Future studies will be directed towards achieving this critical goal.

## METHODS

### Cell lines and reagents

MIAPaCa-2, Panc03.27, CALU-1, H23, H358, H2030, H1373, SW1573, and H460 cells were from laboratory stocks, obtained as described (35). H1792 and H2122 cells were obtained from Dr. Thales Papagiannakopulos (NYU School of Medicine). NYU 59 primary low-passage human pancreatic cancer PDX-derived cells were from Dr. Diane Simeone (NYU School of Medicine), and were generated as described (35). KPC 1203 cells were the gift of Dr. Dafna Bar-Sagi (NYUSoM), and were derived from a pancreatic tumor in an *LSL-Kras*^G12D^/*Tp53*^R172H/+^; Pdx1-Cre (KPC) mouse on C57BL/6 background, as described (81). Immortalized “RAS-less” (*Nras*−*/*−; *Hras*−*/*−, *Kras*^*f/f*^, CreER^Tam^) mouse embryonic fibroblasts were provided by the NCI RAS Initiative at the FNLCR under an MTA.

All cells were grown in 5% CO_2_ at 37C° under media conditions described by the vendor or the source laboratory; details are available from CF upon request. Cells were tested routinely (every 3 months) for mycoplasma contamination by PCR (82), and genotyped by STR analysis at IDEXX Bioresearch. SHP099 was purchased from Wuxi. ARS1620 and AMG510 were purchased from Selleckchem. MRTX1257 was provided by Mirati Therapeutics under a collaborative agreement.

### Plasmids and virus production

A human SHP2 cDNA was cloned into pMSCV-IRES-GFP, pCW57.1, and PLX304. pCW57.1 and PLX304 were a gift from David Root, Addgene plasmid #41393 and # 25890. Mutations were introduced by using the QuikChange II site-directed mutagenesis kit (Agilent Technologies). Sequences encoding Kras and SHP2 sgRNAs (Supplementary Table x) were cloned into the BbsI site of pX458 (a gift from Feng Zhang, Addgene plasmid # 48138). The pTRIPZ sh*SOS1* construct was a gift from Dr. Dafna Bar-Sagi (NYUSoM).

Viruses were produced by co-transfecting HEK293T cells with lentiviral or retroviral constructs and packaging vectors (pVSV-G + pvPac for retroviruses; pVSV-G + dR8.91 for lentiviruses). Forty-eight (48) hours later, culture media were passed through a 0.45 mm filter, and viral supernatants, supplemented with 8 μg/ml of polybrene (Sigma), were used to infect 70% confluent cells in six-well dishes for 16h at 37 °C. Stable pools were selected either by using the appropriate antibiotic or by fluorescence activated cell sorting (FACS) for EGFP.

### G12C targeting

KPC 1203 cells (2×10^5^) were co-transfected with 2ug of the Cas9/sgRNA vector PX458 and 4 ul of ssODN HDR template (20 uM) using Xtreme gene (Roche). Two days following transfection, GFP+ cells were purified by FACS, and single cells were seeded into a 96-well plate. Clones were screened by immunoblotting with RAS^G12D^-specific Rabbit mAb D8H7 (Cell Signaling #14429). Further characterization of the only clone (1/96) that had lost KRAS^G12D^ expression was performed by analyzing genomic DNA. The region flanking *Kras* exon 1 was amplified by PCR, and the product was sub-cloned into the Zero Blunt TOPO vector (Thermo #K287540), followed by Sanger sequencing of the insert using M13 primers. See Table S3 for sequences.

### RAS nucleotide exchange

Purified RAS proteins used in binding experiments were prepared by diluting stock protein (typically containing 20-250 µM RAS) 25-fold with 20 mM Tris-Cl buffer pH 7.5 containing 5 mM EDTA, 0.1 mM DTT, and 1 mM (final concentration) of nucleotide (GDP or GTPγS). Samples were incubated at 30°C for 30 minutes. MgCl2 was then added to a final concentration of 20 mM, and the solution was incubated on ice for at least 5 minutes prior to use.

### Selection of phage-displayed antibody fragments against 12C-ARS

General procedures for the development of Fabs against purified protein targets have been described (83). Four rounds of phage display library selection with biotinylated KRAS(G12C)-GDP+ARS1620 at 100 nM, 100 nM, 50 nM, and 20 nM were performed. The first round recovered clones that bound to KRAS^G12C^-GDP+ARS1620; the second round recovered clones that bound to KRAS^G12C^-GDP+ARS1620, previously pre-cleared with KRAS^G12C^-GDP; the third round recovered clones that bound to KRAS^G12C^-GDP+ARS1620, previously pre-cleared with KRAS^G12C^-GTP. The final round recovered clones that bound to KRAS^G12C^-GDP+ARS1620, previously precleared with KRAS(G12C)-GDP. Phage captured on beads were eluted in 100 μl of 0.1 M Gly-HCl (pH 2.1) and immediately neutralized with 35 μl of 1M Tris-Cl (pH 8). Recovered clones were analyzed by phage ELISA and DNA sequencing, as described (83).

### Expression, purification and characterization of recombinant Fabs

Phage display vectors were converted into Fab expression vectors that contain a substrate tag for the biotin ligase BirA (AviTag, Avidity, LLC) at the carboxyl terminus of the heavy chain. Fabs were expressed in *E. coli* strain 55244 (ATCC), and were purified by protein G affinity chromatography, followed by cation exchange chromatography, as described (83). Purified Fabs were biotinylated *in vitro* using purified BirA. Approximately 2–5 mg of purified Fabs were obtained routinely from a 1 L bacterial culture. SDS-PAGE showed that Fabs were >90% pure. Fab binding to targets was assessed by a bead binding assay, as described previously (84). Briefly, biotinylated Fabs were immobilized on Dynabeads M280 streptavidin, and then excess biotin-binding sites on streptavidin were blocked with biotin. Biotinylated RAS proteins were titrated into the solution containing Fab-immobilized beads. After washing, biotinylated RAS proteins bound to Fabs on the beads were detected with neutravidin Dylight650. The median signal intensity in the Dylight650 channel for the 75-95th percentile population was taken as representative. The 12C-ARS Fab showed the highest specificity among Fabs tested, so this antibody was used for further analyses. The DNA sequence of 12C-ARS Fab is shown in Table S3.

### Proliferation assays

Cells (500-2,000/well) were seeded into 96-well plates. Following incubation with DMSO, 10 μM ARS1620, 10 μM SHP099 or both drugs, cell viability (n =3) was assayed at different times using the PrestoBlue cytotoxicity assay (Thermo Fisher), according to the manufacturer’s protocol. Media (including drugs) were refreshed every 48 h. Briefly, 10 μL of PrestoBlue reagent were added to each well, and after 2h, fluorescence was measured on a multi-plate reader, with excitation wavelength of 530 nm/emission wavelength of 590 nm. Data were corrected for PrestoBlue background fluorescence in media alone. All data represent at least two biological independent experiments in which technical triplicates were performed.

### Clonogenic survival assays

Cells (100-2,000) were seeded in six-well plates one day before treatment with DMSO, 10 μM ARS1620, 10 μM SH099 or both drugs, allowed to grow until they formed colonies (7-14 days), rinsed twice with PBS to remove floating cells, fixed in 4% formaldehyde in PBS (v/v) for 10-15 minutes, and stained in 0.1% crystal violet/10% ethanol for 20 minutes. Staining solution was aspirated, and colonies were washed with water 3x, air-dried and visualized with an Odyssey Imaging System (LICOR). Results were quantified by using the ImageJ Colony Area PlugIn (85). At least three biological replicates were performed.

### Immunoblotting

Whole cell lysates were generated in modified radioimmunoprecipitation (RIPA) buffer (50mM Tris-HCl pH 8.0, 150mM NaCl, 2mM EDTA, 1% NP-40, and 0.1% SDS, without sodium deoxycholate), supplemented with protease (40µg/ml PMSF, 2µg/ml antipain, 2µg/ml pepstatin A, 20µg/ml leupeptin, and 20µg/ml aprotinin) and phosphatase (10mM NaF, 1mM Na_3_VO_4_, 10mM β-glycerophosphate, and 10mM sodium pyrophosphate) inhibitors. After clarification of debris by a microfuge, samples were quantified with the DC Protein Assay Kit (Bio-Rad). Total lysate protein was resolved by standard SDS-PAGE and transferred in 1X transfer buffer and 15% methanol. Membranes were incubated with their respective primary and secondary antibodies labeled with IRDye (680nm and 800nm) and then visualized using the LICOR. Antibodies against phospho-p42/44 MAPK (rabbit polyclonal; #9101; 1:1000) were obtained from Cell Signaling. Moncolonal pan-RAS antibody (clone Ab-3; OP40-100UG; 1:1000) was obtained from Millipore. Rabbit polyclonal antibodies against SHP2 (sc-280; 1:1000) and mouse monoclonal ERK-2 (D2: sc-1647; 1:1000) were purchased from Santa Cruz Biotechnology. Mouse monoclonal anti-SOS1 (# MA5-17234) was purchased from Invitrogen. Rabbit monoclonal anti-DUSP6 antibody EPR129Y was obtained from Abcam (# ab76310).

### KRAS-G12C activity measurements

Cells cultured in 6-well plates were treated as described in the Figures with G12C-I (ARS1620, AMG510, and/or SHP099). Cells were lysed by incubating them in GTPase lysis buffer (25 mM Tris-Cl pH7.2, 150 mM NaCl, 5 mM MgCl2, 1% NP-40 and 5% glycerol supplemented with protease inhibitors and phosphatase inhibitors on ice for 15 minutes immediately before analysis. After centrifugation for 15 minutes at 15,000g, supernatants were collected and incubated with streptavidin (SA) agarose resin (Thermo Fisher Scientific) for 1 hour at 4°C, followed by a brief centrifugation, to decrease non-specific binding to the resin. Pre-cleared lysates were incubated with biotinylated 12C/V-MB or 12C-ARS-Fab bound to SA agarose for 1.5 hours at 4°C while rotating. Agarose beads were then washed twice with GTPase lysis buffer, boiled in 1x SDS-PAGE sample buffer, subjected to immunoblotting with a pan-RAS antibody (Millipore).

### LC/MS-MS Assay for ARS binding to KRAS^G12C^

Cells (5 x10^5^) were treated with the indicated compounds for the times listed and subsequently washed twice with PBS and prepared for protein extraction and LC/MS-MS analysis, as described (15). LC/MS-MS was performed at the PCC Proteomics Shared Resource at NYU School of Medicine.

### Histology and Immunohistochemistry

H&E, Masson-Trichome staining, and immunohistochemistry (IHC) were performed by the PCC Experimental Pathology Shared Resource at NYU School of Medicine. IHC for pERK (Cell Signaling, 4370), CD31 (Cell Signaling, D8V9E), Cleaved Caspase 3 (Cell Signaling, D3E9), Ki67 (Spring Biosciences, SP6), αSMA (Abcam, ab5694) was performed on sections from paraformaldehyde-fixed tumors.

### Xenografts

All animal experiments were approved by, and conducted in accordance with the procedures of, the IACUC at New York University School of Medicine (Protocol no.170602). NY53 and H2122 xenografts were established by sub-cutaneous injection of 5 × 10^6^ cells in 50% Matrigel (Corning) into the right flanks of 8-10-week-old *nu/nu* mice (#088 Charles River). Each treatment group contained 8-10 mice. When tumors reached 100-500 mm^3^, as measured by calipers (size=length*width^2^*0.5), mice were randomized to four groups (10 mice/group) for each model, and treated with: (i) vehicle, (ii) SHP099, (iii) ARS1620, or (iv) SHP099/ARS1620. Investigators were not blinded to group allocation. The following oral gavage dosing regimens were employed: ARS1620 (200mg/kg QD), SHP099 (75mg/kg QD), or ARS1620 200 mg/Kg QD, SHP099 75 mg/kg QD. SHP099 was resuspended in 0.6% methylcellulose, 0.5% Tween80 in 0.9% saline. ARS1620 was dissolved 1% N-methyl-2-pyrrolidone (NMP) + 19% polyethylene glycol 400 (PEG400) + 80% (10% hydroxypropyl in water). Caliper and weight measurements were performed every other day and continued until termination of the experiments.

### Orthotopic PDAC model

KCP cells were generated as described above. Cells (1 × 10^5^) were suspended in Matrigel, implanted into the pancreata of 6-8-week-old syngeneic mice, and allowed to establish for 14-30 days before beginning treatment. Vehicle, ARS1620 (200mg/kg QD), SHP099 (75mg/kg QD), or ARS1620 200 mg/Kg QD, SHP099 75 mg/kg QD was administered for the indicated time, and mice were euthanized. Where indicated, α-PD-1 antibody (200 μg; RMP1-14, BioXcell) was used. Dosing was repeated every three days for the duration of the experiment. Control mice were injected with PBS or isotype control antibody (clone LTF-2, BioXcel).

### NSCLC GEMM studies

*KRAS*^*G12C*^ or *KRAS*^*G12C*^;*Tp53*^*R270H*^ mice (67) were monitored by MRI for tumor development after intranasal induction with adeno-Cre (2.5 × 10^6^ pfu). Tumor-bearing mice were dosed with Vehicle, ARS1620 (200mg/kg QD), SHP099 (75mg/kg QD), or ARS1620 200 mg/Kg QD, SHP099 75 mg/kg QD and monitored by MRI every 2 weeks. In some experiments, MRTX1257 (50 mg/Kg QD) was used in place of ARS.

### Flow Cytometry

Tumors were minced, chopped, and digested in DMEM containing 2.0 mg/mL collagenase IV (GIBCO), 1.0 mg/mL hyaluronidase (Worthington), 0.1% soybean trypsin inhibitor, 50U/mL DNase I (STEMCELL Technologies) at 37°C for 1h. Single cell suspensions were obtained by passage through a strainer (70 µm), washed in FACS buffer (PBS with 5% FBS), incubated with LIVE/DEAD Fixable Zombie Yellow Fixable Viability Kit (Biolegend, 423104) for 30 min., blocked with anti-CD16/32 (Biolegend, clone 93) for 5 min. on ice, and then incubated with fluorophore-conjugated antibodies on ice for 45 min. For detection of intracellular markers, FOXP3 Fixation/Permeabilization Buffer Set (BioLegend) was used, according to the manufacturer’s instructions. Antibodies for flow cytometry are described in Table S4. For quantifying apoptosis, cells were stained by using the PE Annexin V Apoptosis Detection Kit I according to manufacturer’s protocol (BD). Flow cytometry was performed on an LSR II flow cytometer (BD) at the PCC Precision Immunology Shared Resource at NYU School of Medicine and analyzed by using FlowJo software (BD).

### RNA Extraction and Sequencing

RNA was extracted from frozen tumors using the miRNeasy Mini Kit (Qiagen), according to the manufacturer’s instructions. RNA sequencing was performed by the PCC Genome Technology Center Shared Resource (GTC). Libraries were prepared by using the Illumina TruSeq Stranded Total RNA Sample Preparation Kit and sequenced on an Illumina NovaSeq 6000 using 150 bp paired-end reads. Sequencing results were de-multiplexed and converted to FASTQ format using Illumina bcl2fastq software. The average number of read pairs/sample was 35.4 million. Data were processed by the Perlmutter Cancer Center Applied Bioinformatics Laboratories shared resource (ABL).

### qRT-PCR

Total cellular RNA was isolated by using the Qiagen RNeasy kit. cDNA was generated by using the SuperScript IV First Strand Synthesis System (Invitrogen) for RT-PCR. qRT-PCR was performed with Fast SYBR™ Green Master Mix (Applied Biosystems), following the manufacturer’s protocol, in 384-well format in C1000 Touch Thermal Cycler (Biorad). Differential gene expression analysis was performed with CFX Manager (Biorad) and normalized to GAPDH expression. Primers used are listed in Table S3.

### Bliss Analysis

Potential drug synergy was assessed by Bliss analysis as: *Y*_*ab,P*_ = *Y*_*a*_ + *Y*_*b*_ – *Y*_*a*_*Y*_*b*_, where *Y*_*a*_ stands for percentage inhibition of drug *a* and *Y*_*b*_ stands for percentage inhibition of drug *b* (86). Synergistic effects were defined as % of observed effect greater than *Y*_*ab,P*_.

### Statistical Analysis

Data are expressed as mean ± standard deviation. Significance was assessed using Welch’s t test, or one-way ANOVA, as appropriate. Survival rates were analyzed by log-rank test. Statistical analyses were performed in Prism 8 (GraphPad Software). Significance was set at P = 0.05.

## Data availability

RNA sequence data have been deposited in the GEO database under the accession code GSE149815. All other data supporting the findings of this study are available within the article, the Supplemental information files, or from the corresponding author upon request.

## Author contributions

Conception and design: C. Fedele, K-K. Wong, B.G. Neel; Acquisition of data: C. Fedele, S. Li, K. W. Teng, C. Foster, D. Peng, H. Ran, T. Hattori, A. Koide, Y. Wang, J. Leinwand, W. Wang, B. Diskin; Analysis and interpretation of data: C. Fedele, S. Li, K. W. Teng, C. Foster, D. Peng, H. Ran, T. Hattori, A. Koide, H. Ran, P. Mita, M. Geer, K.H. Tang, I. Dolgalev, U. Ozerdem, G. Miller, S. Koide, K.-K. Wong, B. G. Neel; Administrative, technical, or material support: C. Fedele, S. Li, W. Teng, C. Foster, D. Peng, H. Ran, T. Hattori, A. Koide, H. Ran, P. Mita, M. Geer, K.H. Tang, I. Dolgalev, U. Ozerdem, J. Deng, T. Chen, J. Leinwand, W. Wang, B. Diskin; Writing the manuscript: C. Fedele, B.G. Neel; Study supervision: C. Fedele, K-K. Wong, B.G. Neel.

## Acknowledgements

We thank Drs. Thales Papagiannakopoulos and Diane Simeone for sharing cell lines, Drs. Dafna Bar-Sagi, David Root, and Feng Zhang for plasmids, and the PCC Experimental Pathology, Precision Immunology, Genome Technology Center, Applied Bioinformatics Laboratories (ABL) and Proteomics Laboratory shared resources (P30CA016087) for technical support. We thank Dr. Jamie Christensen (Mirati Therapuetics) for generously providing MRTX 1275 and for helpful discussions. We also thank Drs. Toshiyuki Araki, Kiyomi Araki, Abhishek Bhardwaj, Jayu Jen, and Shuang Zhang and Ms. Angel Sing and Wei Wei (Neel Lab) for advice and discussion on this project. This work was supported by CA49152 (B.G.N), CA219670 (K.K.W), CA194864 (S.K.) and Cancer Center Core Grant P30 CA016087.

