## supplemental data for "SHP2 Inhibition Abrogates Adaptive Resistance to KRAS^G12C^-Inhibition and Remodels the Tumor Microenvironment of *KRAS*-Mutant Tumors"

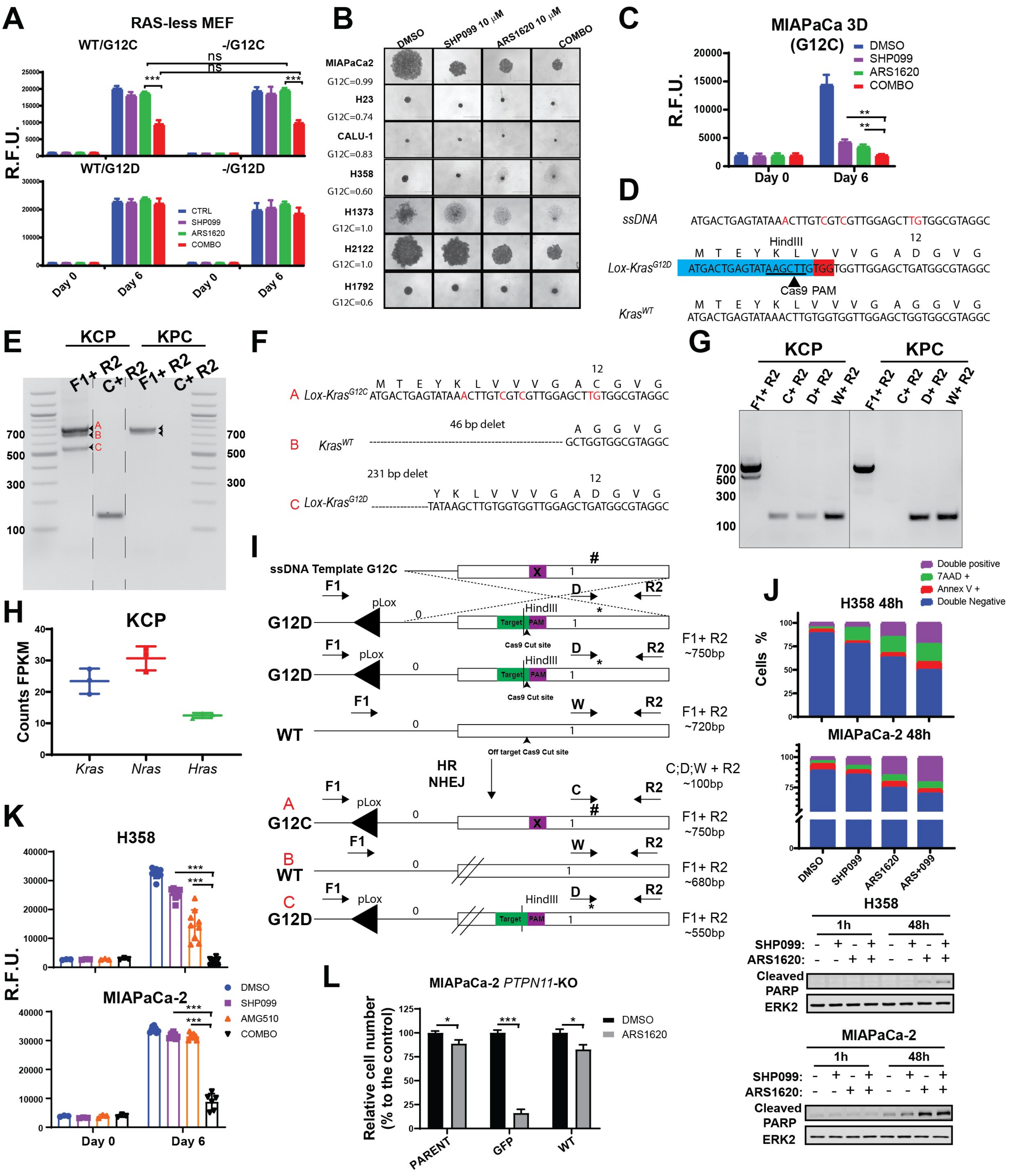

A
RAS-less MEF
WT/G12C
-/G12C
WT/G12D
-/G12D
R.F.U.
ns
***
CTRL
SHP099
ARS1620
COMBO
Day 0
Day 6
B
DMSO
SHP099 10 μM
ARS1620 10 μM
COMBO
MIAPaCa2
G12C=0.99
H23
G12C=0.74
CALU-1
G12C=0.83
H358
G12C=0.60
H1373
G12C=1.0
H2122
G12C=1.0
H1792
G12C=0.6
C
MIAPaCa 3D
(G12C)
DMSO
SHP099
ARS1620
COMBO
R.F.U.
**
Day 0
Day 6
D
ssDNA
ATGACTGAGTATAAACTTGTCGTCGTTGGAGCTTGTGGCGTAGGC
HindIII
12
M T E Y K L V V V G A D G V G
Lox-Kras^G12D
ATGACTGAGTATAAAGCTTGTGGTGGTTGGAGCTGATGGCGTAGGC
Cas9 PAM
Kras^WT
M T E Y K L V V V G A G G V G
ATGACTGAGTATAAACTTGTGGTGGTTGGAGCTGGTGGCGTAGGC
E
KCP
KPC
F1+ R2
C+ R2
700
500
300
100
A
B
C
F
12
M T E Y K L V V V G A C G V G
A Lox-Kras^G12C ATGACTGAGTATAAACTTGTCGTCGTTGGAGCTTGTGGCGTAGGC
46 bp delet
A G G V G
B Kras^WT GCTGGTGGCGTAGGC
231 bp delet
12
Y K L V V V G A D G V G
C Lox-Kras^G12D TATAAGCTTGTGGTGGTTGGAGCTGATGGCGTAGGC
G
KCP
KPC
F1+ R2
C+ R2
D+ R2
W+ R2
700
500
300
100
H
KCP
Counts FPKM
Kras
Nras
Hras
I
ssDNA Template G12C
X
1
#
F1
D
R2
pLox
0
G12D
HindIII
Target
PAM
Cas9 Cut site
*
F1+ R2
~750bp
WT
W
~720bp
Off target Cas9 Cut site
HR
NHEJ
C;D;W + R2
~100bp
A
G12C
C
B
~680bp
~550bp
J
Double positive
7AAD +
Annex V +
Double Negative
H358 48h
MIAPaCa-2 48h
Cells %
DMSO
SHP099
ARS1620
ARS+099
H358
1h
48h
SHP099:
ARS1620:
Cleaved PARP
ERK2
MIAPaCa-2
K
H358
MIAPaCa-2
R.F.U.
DMSO
SHP099
AMG510
COMBO
***
Day 0
Day 6
L
MIAPaCa-2 PTPN11-KO
Relative cell number (% to the control)
DMSO
ARS1620
*
***
PARENT
GFP
WT


FIG. S1

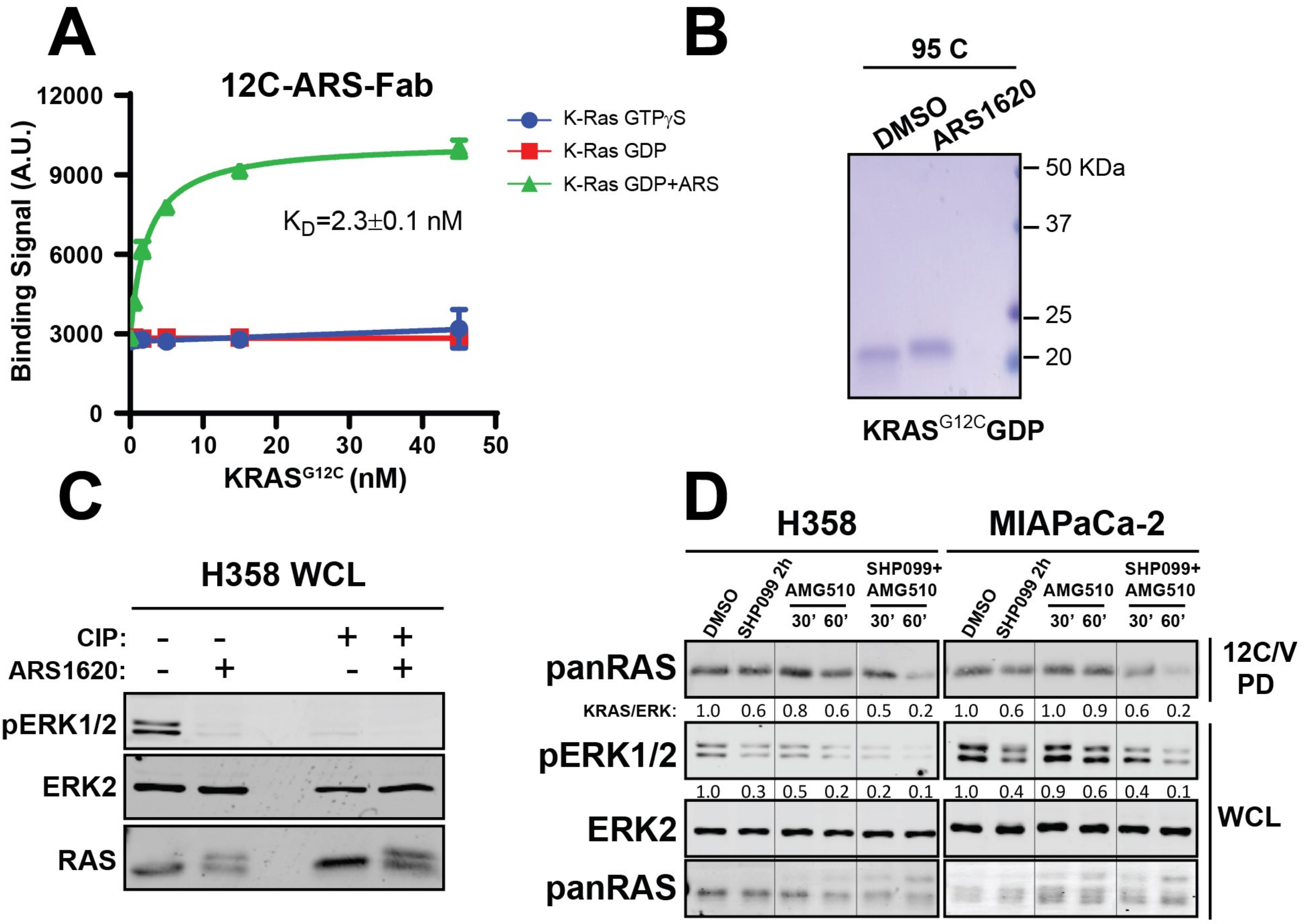
A
12C-ARS-Fab
K-Ras GTPγS
K-Ras GDP
K-Ras GDP+ARS
KD=2.3±0.1 nM
Binding Signal (A.U.)
KRAS G12C (nM)
B
95 C
DMSO
ARS1620
50 KDa
37
25
20
KRAS G12C GDP
C
H358 WCL
CIP: - - + +
ARS1620: - + - +
pERK1/2
ERK2
RAS
D
H358
MIAPaCa-2
DMSO
SHP099 2h
AMG510
SHP099+ AMG510
30' 60'
panRAS
KRAS/ERK: 1.0 0.6 0.8 0.6 0.5 0.2 1.0 0.6 1.0 0.9 0.6 0.2
pERK1/2
1.0 0.3 0.5 0.2 0.2 0.1 1.0 0.4 0.9 0.6 0.4 0.1
ERK2
panRAS
12C/V PD
WCL

FIG. S2

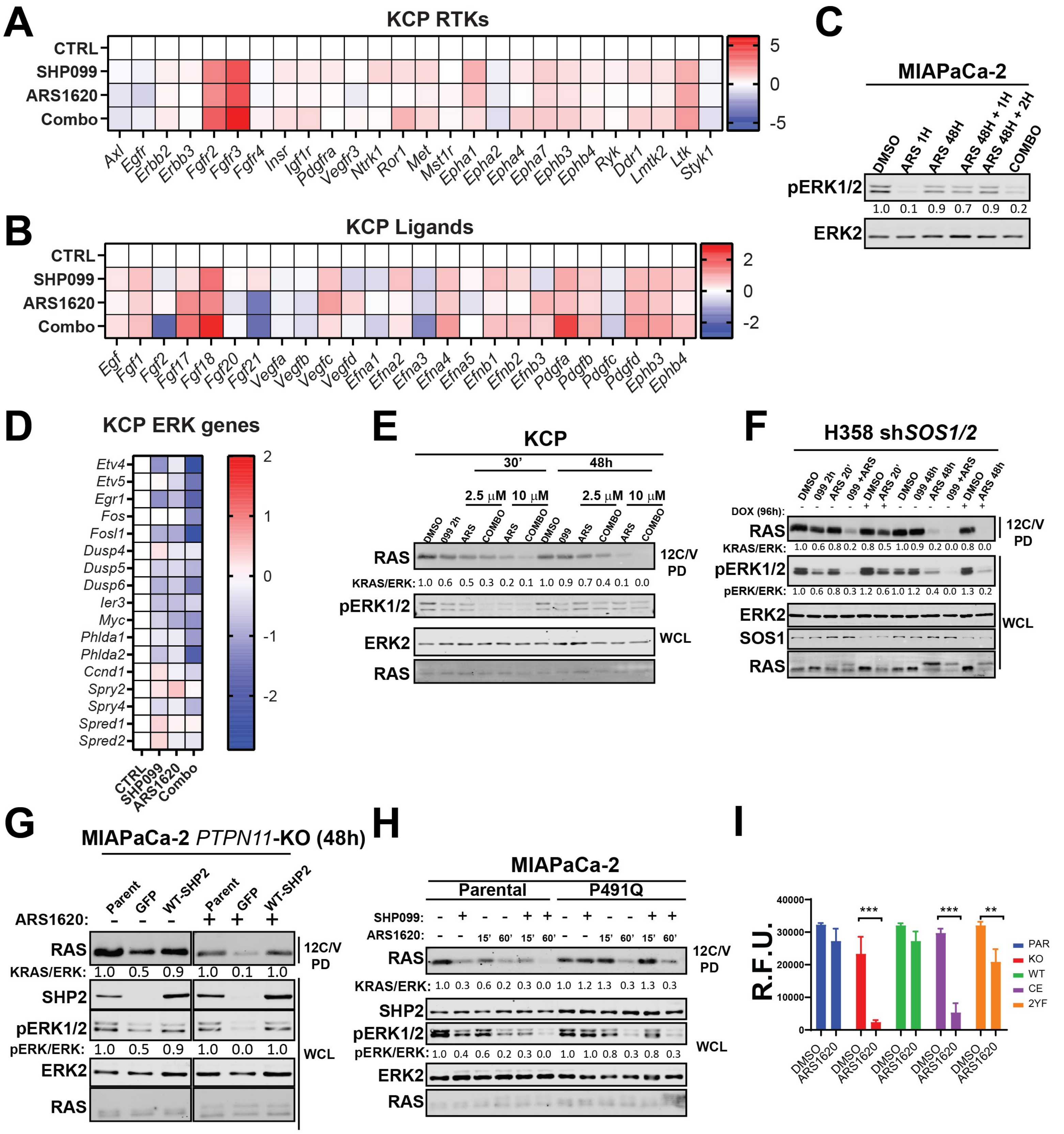
A
KCP RTKs
CTRL
SHP099
ARS1620
Combo
Axl Egfr Erbb2 Erbb3 Fgfr2 Fgfr3 Fgfr4 Insr Igf1r Pdgfra Vegfr3 Ntrk1 Ror1 Met Mst1r Epha1 Epha2 Epha4 Epha7 Ephb3 Ephb4 Ryk Ddr1 Lmtk2 Ltk Styk1
B
KCP Ligands
Egf Fgf1 Fgf2 Fgf17 Fgf18 Fgf20 Fgf21 Vegfa Vegfb Vegfc Vegfd Efna1 Efna2 Efna3 Efna4 Efna5 Efnb1 Efnb2 Efnb3 Pdgfa Pdgfb Pdgfc Pdgfd Ephb3 Ephb4
C
MIAPaCa-2
DMSO ARS 1H ARS 48H ARS 48H + 1H ARS 48H + 2H COMBO
pERK1/2
1.0 0.1 0.9 0.7 0.9 0.2
ERK2
D
KCP ERK genes
Etv4 Etv5 Egr1 Fos Fosl1 Dusp4 Dusp5 Dusp6 Ier3 Myc Phlda1 Phlda2 Ccnd1 Spry2 Spry4 Spred1 Spred2
CTRL SHP099 ARS1620 Combo
E
KCP
30' 48h
2.5 μM 10 μM 2.5 μM 10 μM
DMSO 099 2h ARS COMBO ARS COMBO DMSO 099 ARS COMBO ARS COMBO
RAS
KRAS/ERK: 1.0 0.6 0.5 0.3 0.2 0.1 1.0 0.9 0.7 0.4 0.1 0.0
12C/V PD
pERK1/2
ERK2
RAS
WCL
F
H358 shSOS1/2
DMSO 099 2h ARS 20' 099 +ARS DMSO ARS 20' DMSO 099 48h ARS 48h 099 +ARS DMSO ARS 48h
DOX (96h): - - - - + + - - - - + +
RAS
KRAS/ERK: 1.0 0.6 0.8 0.2 0.8 0.5 1.0 0.9 0.2 0.0 0.8 0.0
12C/V PD
pERK1/2
pERK/ERK: 1.0 0.6 0.8 0.3 1.2 0.6 1.0 1.2 0.4 0.0 1.3 0.2
ERK2
SOS1
RAS
WCL
G
MIAPaCa-2 PTPN11-KO (48h)
Parent GFP WT-SHP2 Parent GFP WT-SHP2
ARS1620: - - - + + +
RAS
KRAS/ERK: 1.0 0.5 0.9 1.0 0.1 1.0
12C/V PD
SHP2
pERK1/2
pERK/ERK: 1.0 0.5 0.9 1.0 0.0 1.0
ERK2
RAS
WCL
H
MIAPaCa-2
Parental P491Q
SHP099: - + - - + + - + - - + +
ARS1620: - - 15' 60' 15' 60' - - 15' 60' 15' 60'
RAS
KRAS/ERK: 1.0 0.3 0.6 0.2 0.3 0.0 1.0 1.2 1.3 0.3 1.3 0.3
12C/V PD
SHP2
pERK1/2
pERK/ERK: 1.0 0.4 0.6 0.2 0.3 0.0 1.0 1.0 0.8 0.3 0.8 0.3
ERK2
RAS
WCL
I
R.F.U.
40000 30000 20000 10000 0
*** *** **
DMSO ARS1620
PAR KO WT CE 2YF

FIG. S3

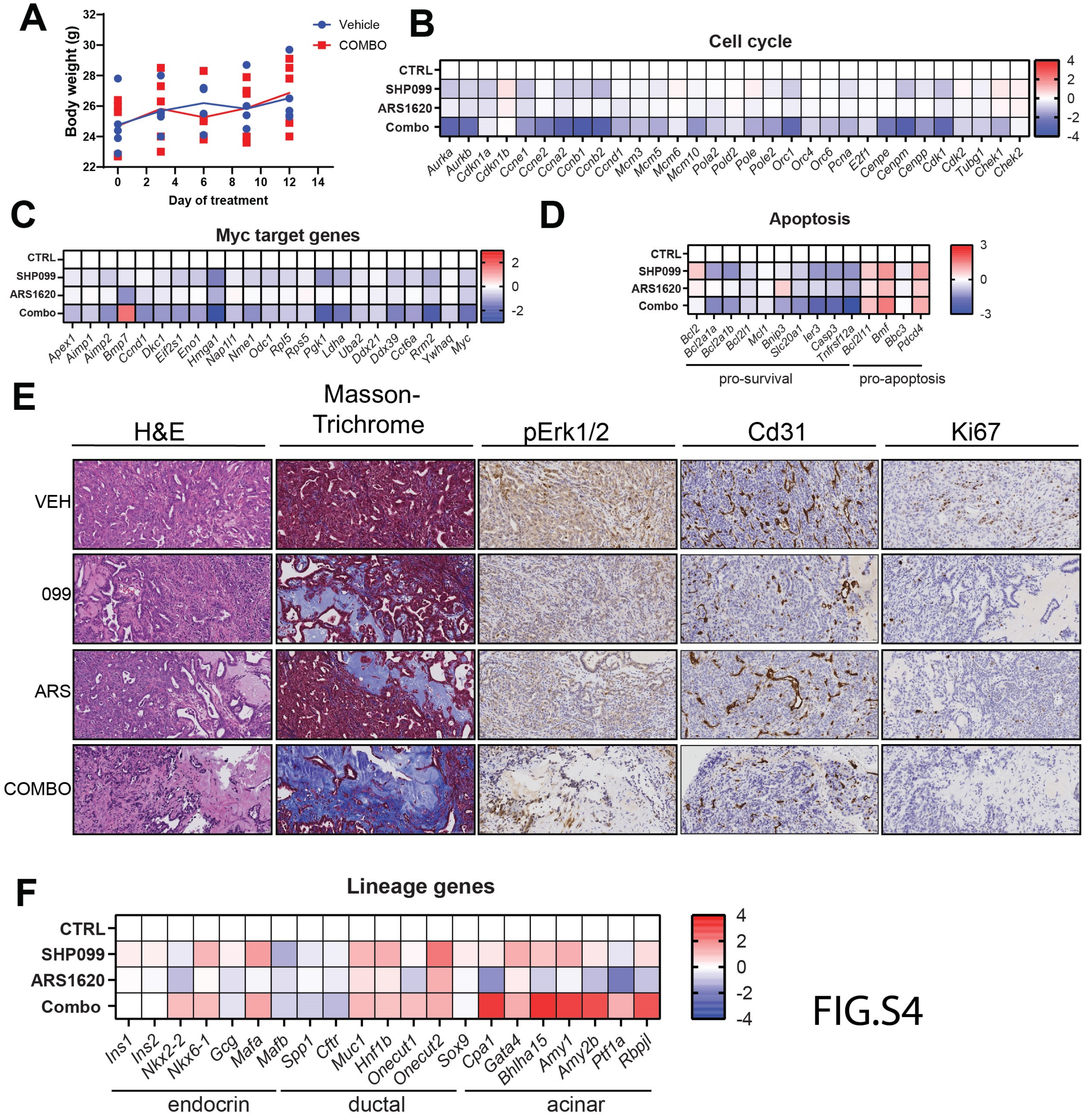

A
Body weight (g)
Day of treatment
Vehicle
COMBO
B
Cell cycle
CTRL
SHP099
ARS1620
Combo
Aurka Aurkb Cdkn1a Cdkn1b Ccne1 Ccne2 Ccna2 Ccnb1 Ccnb2 Ccnd1 Mcm3 Mcm5 Mcm6 Mcm10 Pola2 Pold2 Pole Pole2 Orc1 Orc4 Orc6 Pcna E2f1 Cenpe Cenpm Cenpp Cdk1 Cdk2 Tubg1 Chek1 Chek2
C
Myc target genes
Apex1 Aimp1 Aimp2 Bmp7 Ccnd1 Dkc1 Eif2s1 Eno1 Hmga1 Nap1l1 Nme1 Odc1 Rpl5 Rps5 Pgk1 Ldha Uba2 Ddx21 Ddx39 Cct6a Rrm2 Ywhaq Myc
D
Apoptosis
Bcl2 Bcl2a1a Bcl2a1b Bcl2l1 Mcl1 Bnip3 Slc20a1 Ier3 Casp3 Tnfrsf12a Bcl2l11 Bmf Bbc3 Pdcd4
pro-survival
pro-apoptosis
E
H&E
Masson-Trichrome
pErk1/2
Cd31
Ki67
VEH
099
ARS
COMBO
F
Lineage genes
Ins1 Ins2 Nkx2-2 Nkx6-1 Gcg Mafa Mafb Spp1 Cftr Muc1 Hnf1b Onecut1 Onecut2 Sox9 Cpa1 Gata4 Bhlha15 Amy1 Amy2b Ptf1a Rbpjl
endocrin
ductal
acinar


FIG.S4

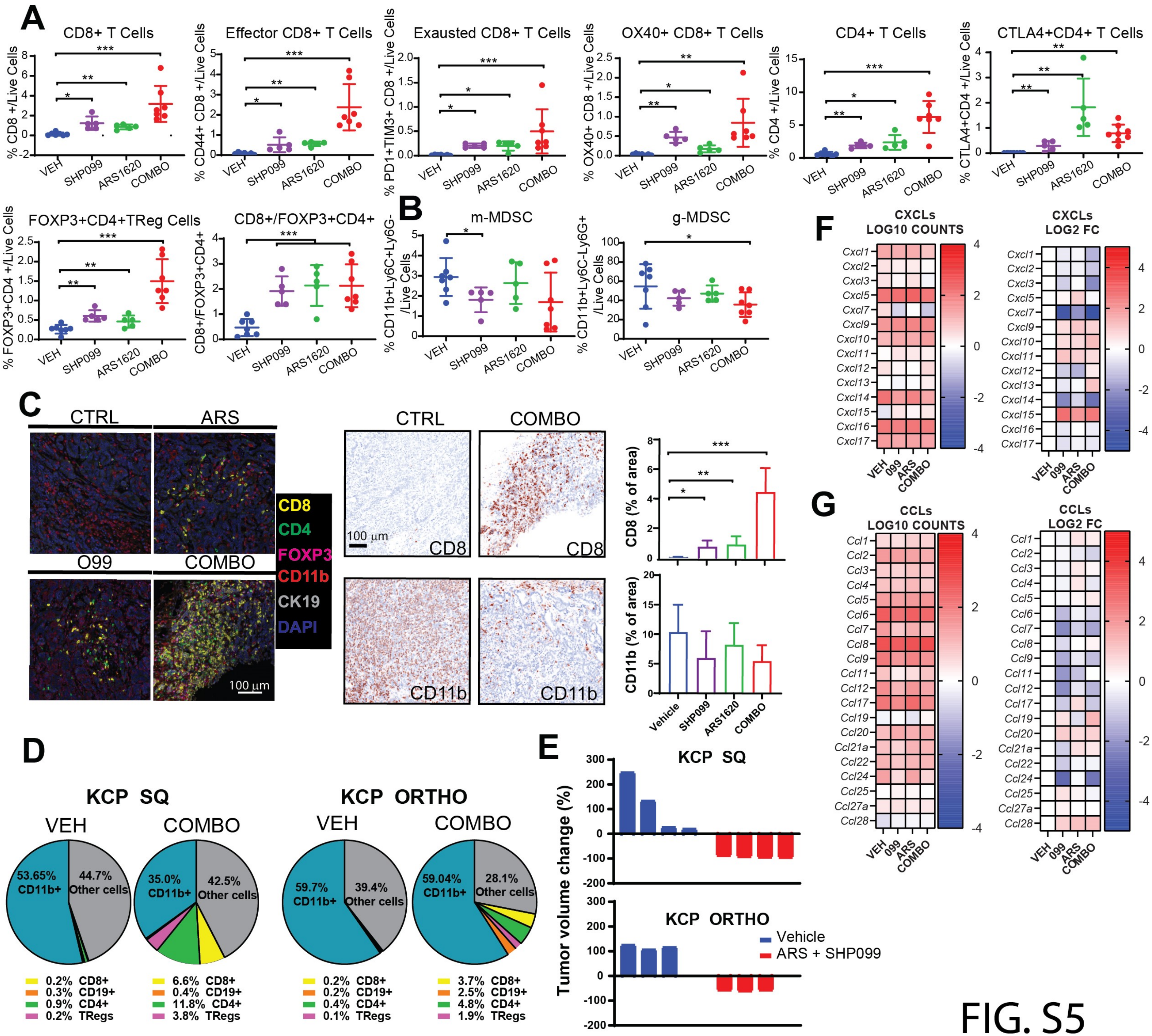

A
CD8+ T Cells
Effector CD8+ T Cells
Exausted CD8+ T Cells
OX40+ CD8+ T Cells
CD4+ T Cells
CTLA4+CD4+ T Cells
FOXP3+CD4+TReg Cells
CD8+/FOXP3+CD4+
VEH
SHP099
ARS1620
COMBO
B
m-MDSC
g-MDSC
C
CTRL
ARS
O99
COMBO
CD8
CD4
FOXP3
CD11b
CK19
DAPI
100 µm
CD8 (% of area)
CD11b (% of area)
Vehicle
D
KCP SQ
KCP ORTHO
53.65% CD11b+
44.7% Other cells
35.0% CD11b+
42.5% Other cells
59.7% CD11b+
39.4% Other cells
59.04% CD11b+
28.1% Other cells
0.2% CD8+
0.3% CD19+
0.9% CD4+
0.2% TRegs
6.6% CD8+
0.4% CD19+
11.8% CD4+
3.8% TRegs
0.2% CD8+
0.2% CD19+
0.4% CD4+
0.1% TRegs
3.7% CD8+
2.5% CD19+
4.8% CD4+
1.9% TRegs
E
Tumor volume change (%)
Vehicle
ARS + SHP099
F
CXCLs LOG10 COUNTS
CXCLs LOG2 FC
G
CCLs LOG10 COUNTS
CCLs LOG2 FC


FIG. S5

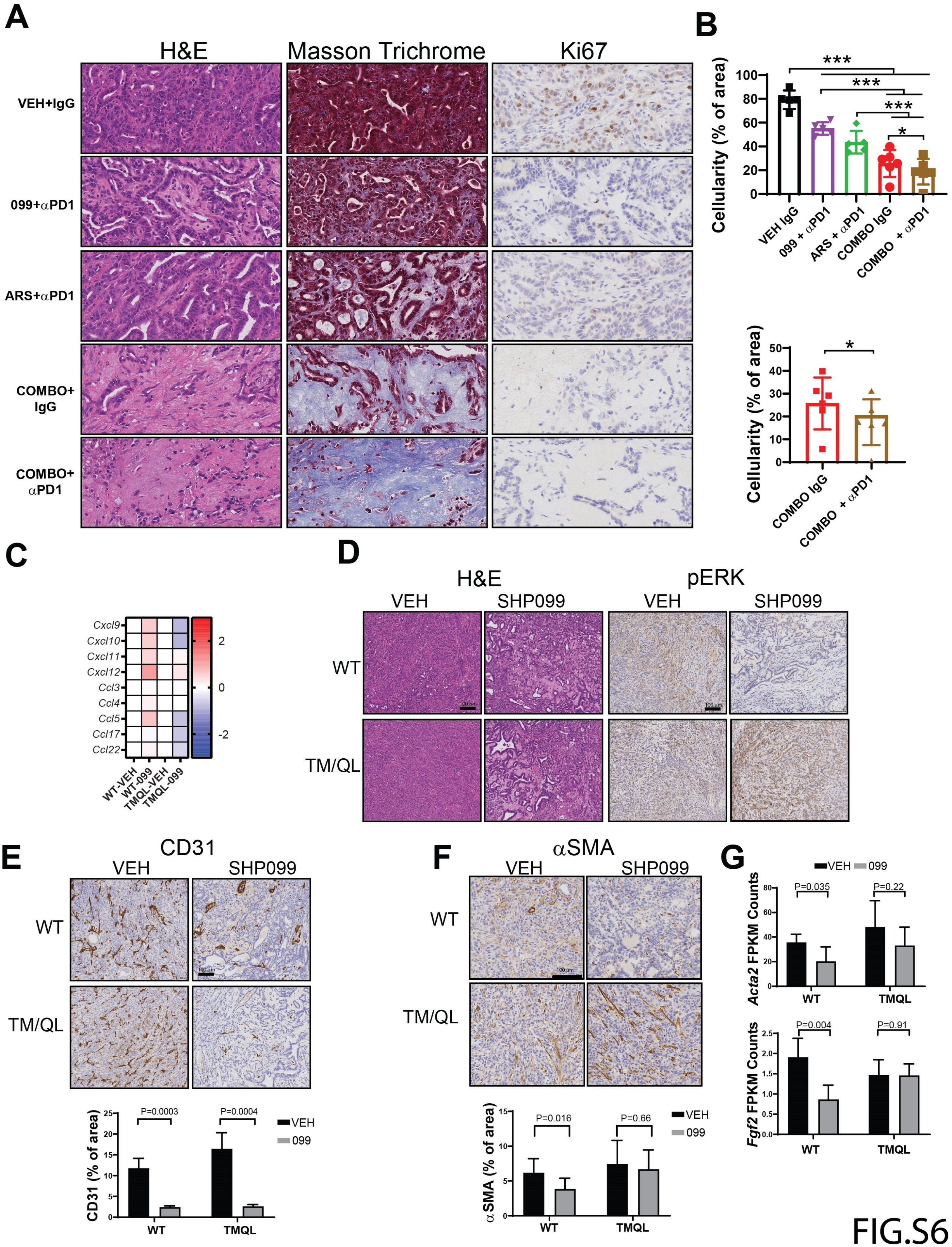

A
H&E
Masson Trichrome
Ki67
VEH+IgG
099+αPD1
ARS+αPD1
COMBO+ IgG
COMBO+ αPD1
B
Cellularity (% of area)
VEH IgG
099 + αPD1
ARS + αPD1
COMBO IgG
COMBO + αPD1
***
*
C
Cxcl9
Cxcl10
Cxcl11
Cxcl12
Ccl3
Ccl4
Ccl5
Ccl17
Ccl22
WT-VEH
WT-099
TMQL-VEH
TMQL-099
D
H&E
pERK
VEH
SHP099
WT
TM/QL
E
CD31
CD31 (% of area)
P=0.0003
P=0.0004
VEH
099
F
αSMA
α SMA (% of area)
P=0.016
P=0.66
G
Acta2 FPKM Counts
P=0.035
P=0.22
Fgf2 FPKM Counts
P=0.004
P=0.91
TMQL


FIG.S6

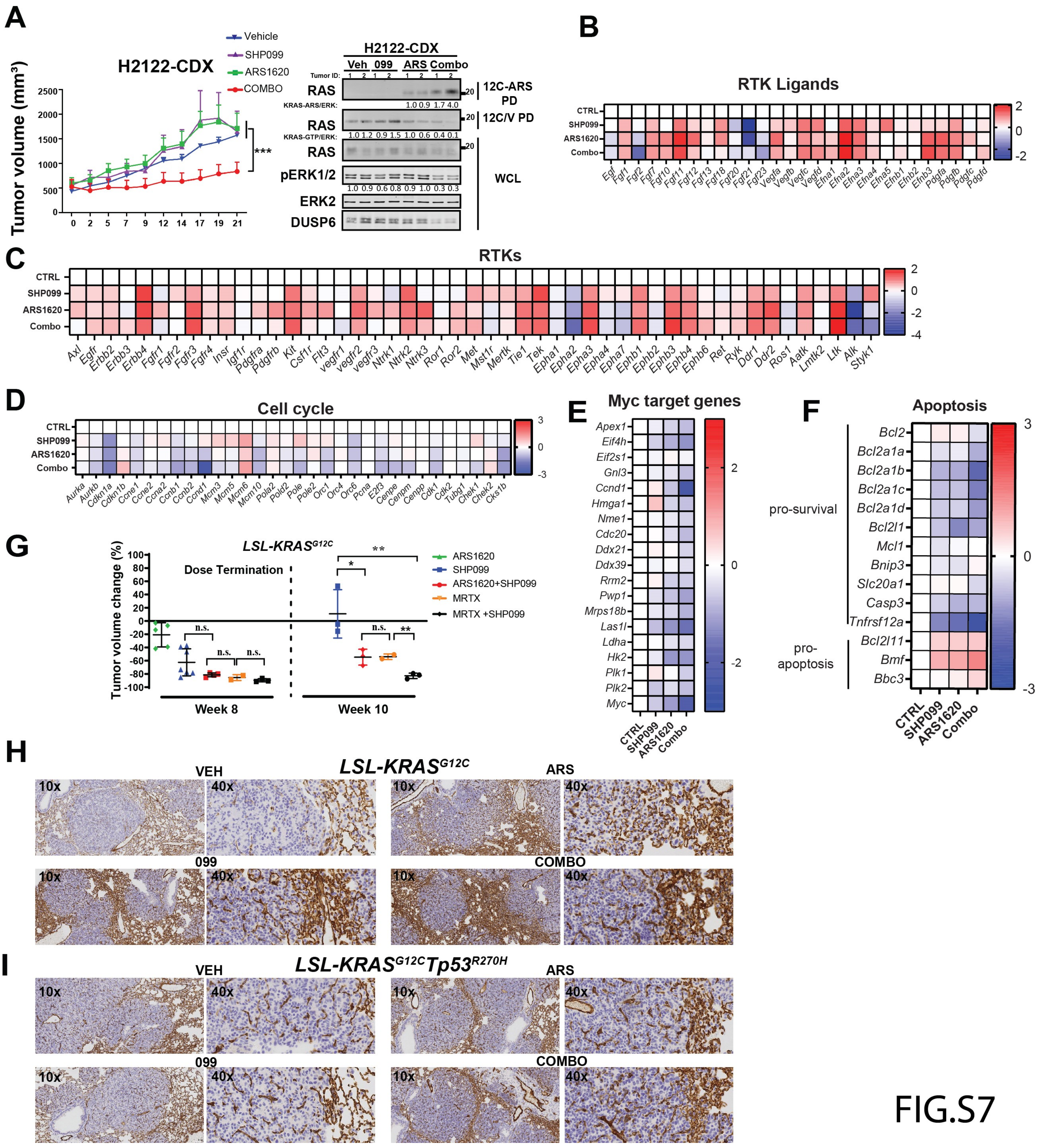

A
H2122-CDX
Vehicle
SHP099
ARS1620
COMBO
Tumor volume (mm³)
***
H2122-CDX
Veh 099 ARS Combo
RAS
12C-ARS PD
RAS
12C/V PD
RAS
pERK1/2
ERK2
DUSP6
WCL
B
RTK Ligands
C
RTKs
D
Cell cycle
E
Myc target genes
F
Apoptosis
pro-survival
pro-apoptosis
G
LSL-KRASG12C
Dose Termination
Tumor volume change (%)
Week 8
Week 10
ARS1620
SHP099
ARS1620+SHP099
MRTX
MRTX +SHP099
H
LSL-KRASG12C
VEH
ARS
099
COMBO
I
LSL-KRASG12CTp53R270H
VEH
ARS
099
COMBO


FIG.S7

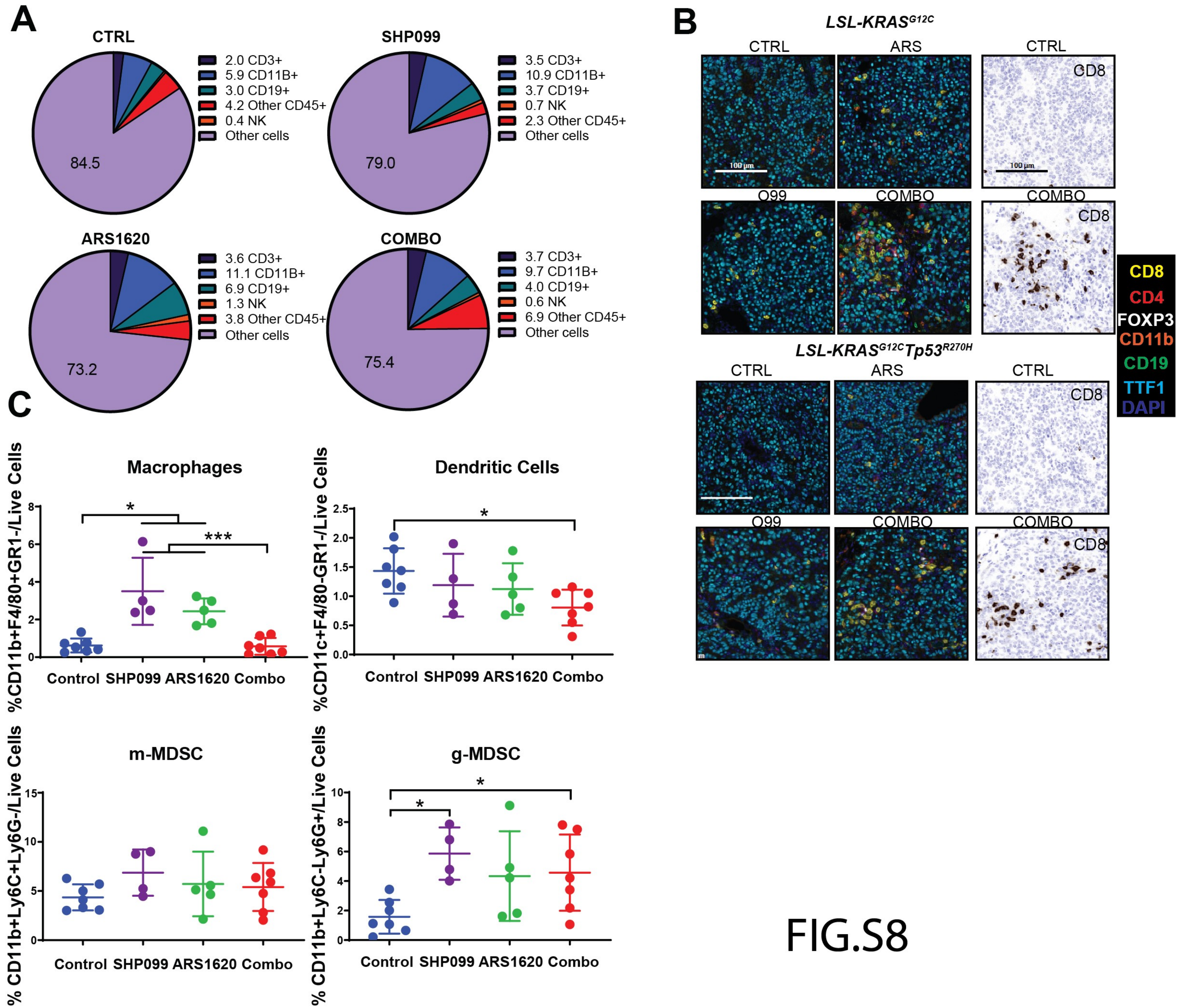
A
CTRL
2.0 CD3+
5.9 CD11B+
3.0 CD19+
4.2 Other CD45+
0.4 NK
Other cells
84.5
SHP099
3.5 CD3+
10.9 CD11B+
3.7 CD19+
0.7 NK
2.3 Other CD45+
Other cells
79.0
ARS1620
3.6 CD3+
11.1 CD11B+
6.9 CD19+
1.3 NK
3.8 Other CD45+
Other cells
73.2
COMBO
3.7 CD3+
9.7 CD11B+
4.0 CD19+
0.6 NK
6.9 Other CD45+
Other cells
75.4
B
LSL-KRAS G12C
CTRL
ARS
CTRL
CD8
100 µm
O99
COMBO
COMBO
CD8
LSL-KRAS G12C Tp53 R270H
CTRL
ARS
CTRL
CD8
O99
COMBO
COMBO
CD8
CD8
CD4
FOXP3
CD11b
CD19
TTF1
DAPI
C
Macrophages
%CD11b+F4/80+GR1-/Live Cells
Control SHP099 ARS1620 Combo
Dendritic Cells
%CD11c+F4/80-GR1-/Live Cells
Control SHP099 ARS1620 Combo
m-MDSC
% CD11b+Ly6C+Ly6G-/Live Cells
Control SHP099 ARS1620 Combo
g-MDSC
% CD11b+Ly6C-Ly6G+/Live Cells
Control SHP099 ARS1620 Combo

FIG.S8

### SUPPLEMENTARY FIGURE LEGENDS

**Supplementary Figure S1. SHP2 inhibition enhances KRAS$^{G12C}$ inhibitor effects in PDAC and NSCLC cell lines.** Cell viability, by PrestoBlue assay, was assessed at 6 days in *Kras$^{wt}$*/*KRAS$^{G12C}$* and *Kras$^{-/-}$*/*KRAS$^{G12C}$* MEFs. **B,** Micrographs of spheroid cultures of *KRAS$^{G12C}$*-expressing cells, treated as indicated for 6 days. Numbers at left indicate *KRAS$^{G12C}$* allele fraction **C,** Cell viability, assessed by PrestoBlue assay, at 0 and 6 days of MIAPaCa-2 spheroid cultures. **D,** Schematic showing strategy used to generate Lox-*Kras*$^{G12C}$ allele from the pancreatic KCP 1203 cells, which carry a Lox-*Kras$^{G12D}$* allele and a pancreas-specific Cre. KCP 1203 cells were co-transfected with a vector expressing *Cas9* and a *Kras*-targeted sgRNA, together with a single-stranded donor oligonucleotide (ssODN) template bearing the new mutation. **E**, PCR products using F1 +R2 primers to discriminate between mutant-*Kras* (~750bp) and WT-*Kras* (~720bp) and C1+R2 to detect the presence of the new G12C mutation in the KCP (G12C) clone, compared with parental KPC 1203 (G12D) cells. Primer FWD F1 flanks the Lox region in intron 0, primer REV R2 anneals the end of exon 1, FWD C1 specifically anneals on exon 1 in the presence of the new generated G12C mutation. **F**, Sanger sequencing of TOPO-cloned PCR products (A-C in red) from E. **G,** Allele-specific PCR using FWD primers WT (W), G12D (D) and G12C (C) and REV R2 in KCP G12C clone and parental KPC 1203 (G12D) cells. **H,** FKPM counts for *Kras* alleles in KCP cells. **I,** Summary of genetic events that generated the new Lox-*Kras*$^{G12C}$ allele starting from parental KCP 1203 cells. **J,** Cell death, after 48h of drug treatment, quantified by flow cytometry and Annexin V/7AAD staining (top)**,** Immunoblot for cleaved PARP3 in lysates from MIAPaCa-2 and H358 cells, treated as indicated (bottom). **K,** Viability of MIAPaCa-2 and H358 cells, assessed by PrestoBlue assay, after 6 days of treatment with DMSO, SHP099, AMG510 (0.1 µM) or Combo. **L,** PrestoBlue assay (6 days) in parental MIAPaCa-2 cells and *PTPN11*-KO MIAPaCa-2 cells reconstituted with GFP or wild type *PTPN11* (WT). For all experiments, drug doses were: SHP099 10 µm/l, ARS1620 10 µm/l, AMG510 0.1 µm/l. Data represent mean ± SD; *P< 0.05, **P< 0.01, ***P < 0.001, one-way ANOVA with Tukey's multiple comparison test.

**Supplementary Figure S2. SHP099 increases KRAS$^{G12C}$-ARS1620 adducts. A,** 12C-ARS Fab binding to KRAS-G12C with/without ARS and GTPγS or GDP**. B,** Coomassie-stained SDS-PAGE of purified, recombinant KRAS$^{G12C}$, pre-incubated with DMSO or ARS1620 for 2h. **C,** Immunoblots of whole cell lysates (WCL) from H358 cells, treated with ARS1620 for 2h or left untreated, with or without incubation with calf-intestinal phosphatase (CIP). Note that CIP treatment does not affect migration of KRAS, arguing against phosphorylation as the cause of the mobility shift. By contrast, the pERK signal is eliminated by CIP treatment. **D,** Immunoblots of WCL and 12C/V MB pull-downs (PD) from H358 and MIAPaCa-2 cells after treatment with DMSO, SHP099, AMG510, or Combo, as indicated. For all experiments, drug doses were: SHP099 10 µm/l, ARS1620 10 µm/l, AMG510 0.1 µmol/l.

**Supplementary Figure S3. SHP099 acts upstream of RAS to block G12C-I–evoked ERK pathway reactivation. A-B,** Time-dependent increase in RTK (A) and RTK ligand (B) gene expression in KCP cells treated for 48h with DMSO, SHP099, ARS1620, or COMBO, determined by RNAseq (colors indicate log2FC). **C,** ERK reactivation, as shown by immunoblot of lysates from MIAPaCa-2 cells treated with DMSO, ARS1620, or ARS1620 + SHP099 for the indicated times. **D,** ERK-dependent gene expression in KCP cells, assessed by RNAseq. **E-H,** Immunoblots of whole cell lysates and 12C/V MB pull-downs (PD) from KCP cells (E), H358 cells expressing DOX-inducible *SOS1* shRNA (sh*SOS*), plus/minus DOX, as indicated (F), parental MIAPaCa-2 cells, or MIAPaCa-2 cells *PTPN11*-KO expressing GFP or reconstituted with WT-*PTPN11,* treated with ARS1620 for 48 h (G), MIAPaCa-2 cells expressing SHP099-resistant *PTPN11* mutant (P491Q) or wild type PTPN11 (WT) treated as described for the indicated times (H) **I,** PrestoBlue assay, performed on *PTPN11*-KO or *PTPN11* KO MIAPaCa-2 cells reconstituted with WT, C459E (CE), Y542F+Y580F (2YF) *PTPN11,* after 6 days of treatment with ARS1620 or DMSO. For all experiments, drug doses were: SHP099 10 µm/l, ARS1620 10 µm/l. Data represent mean ± SD; Significance was assessed by multiple unpaired Student's *t* test (two-tailed)

**Supplementary Figure S4. ARS1620/SHP099 combination is efficacious in PDAC model *in vivo*. A,** ARS1620/SHP099 regimen is well-tolerated in KCP-derived orthotopic tumors, with no significant decrease in body weight after 12 days of treatment. **B-D,** Cell cycle- MYC target- and apoptosis gene expression in KCP-derived orthotopic tumors after Vehicle, SHP099, ARS1620, and COMBO treatment for 3 days, determined by RNAseq (colors indicate log2FC) **E,** H&E, Masson Trichrome, CD31, pERK, and Ki67 staining in KCP tumors treated for 10d with Vehicle, SHP099, ARS1620, of COMBO. 20x magnification, Scale bar =100µm. **F,** Pancreatic epithelial lineage-specific gene expression in Control and treated KCP-derived orthotopic tumors determined by RNAseq (colors indicate log2FC). For all experiments, drug doses were: SHP099 (75 mg/kg body weight, daily), ARS1620 (200 mg/kg body weight, daily), or both drugs (daily).

**Supplementary Figure S5. ARS1620/SHP099 provokes an anti-tumor immune program in syngeneic PDAC model. A-B,** Tumor-infiltrating T cells (A) and MDSCs (B) from KCP-derived orthotopic tumors, analyzed after treatment for 12 days with the indicated drugs. Data represent mean ± SD; **P* < 0.05, ***P* < 0.01, ****P* < 0.001, multiple unpaired Welch's *t* test (two tailed).**C,** Multiplex IF/IHC analysis of tumors from KCP mice, treated as indicated for 12 days, stained with the indicated markers and quantified as shown. Data represent mean ± SD; *P< 0.05, **P< 0.01, ***P< 0.001, one-way ANOVA with Tukey's multiple comparison test. **D,** Pie charts summarizing composition of various immune cells (CD45+) in subcutaneous (SQ) versus orthotopic (ortho) KCP-derived tumors, treated with Vehicle or ARS1620+SHP099 for 10 days. **E,** Changes in volume of subcutaneous (SQ) and orthotopic (ORTHO) KCP tumors established in syngeneic mice and treated with Vehicle or ARS1620+SHP099 for 10 days. Note greater response in SQ tumors. **F-G,** *CXC* (F) and *CCL* (G) chemokine expression in KCP-derived tumors treated for 3 days, assessed by RNAseq (colors are: left, log10 of raw counts averages and log2FC, right).

**Supplementary Figure S6. ARS1620/SHP099 efficacy is enhanced by anti-PD-1 in PDAC model and effect of SHP2 inhibition on PDAC tumor microenvironment. A-B,** H&E, Masson Trichrome and Ki67 staining (A) and quantification (B) from orthotopic KCP tumors, analyzed after treatment for 12 days with the indicated drugs. **C**, Expression of chemokines potentially involved in T cell immigration in tumors from *Ptpn11* KO-KCP cells reconstituted with wild-type (WT) or SHP099-resistant *PTPN11*$^{T253M/Q257L}$ mutant (TM/QL), treated for 10 days with vehicle or SHP099 (75 mg/kg body weight, daily). **D-F,** H&E, pERK, CD31, and $\alpha$SMA staining of representative KCP tumors established as in C. **G,** FKPM counts for *Acta2* (top) and *Fgf2* (bottom) in KCP tumors established as in C. Data represent mean ± SD; **P*< 0.05, **P< 0.01, ***P< 0.001, Student's *t* test (two-tailed).

**Supplementary Figure S7. ARS1620/SHP099 is also efficacious in NSCLC GEMMs. A,** Growth of H2122 cell-derived xenografts (left) and immunoblots (right) of tumor lysates and 12C/V MB or 12C-ARS Fab pull-downs (PD) from mice treated as indicated. **B-F,** Time-dependent expression of RTK (B), RTK ligand (C), cell cycle (D), MYC target (E) and apoptotic (F) genes in *LSL-KRAS*$^{G12C}$-*Tp53*$^{R270H}$ tumors after Vehicle, SHP099, ARS1620, or Combo treatment for 3 days, determined by RNAseq (colors indicate log2FC). **G**, Quantification of tumor volume changes if *LSL*-KRAS$^{G12C}$ NSCLC GEMMs after treatment with vehicle, SHP099 (75 mg/kg, daily), ARS1620 (200 mg/kg, daily), ARS1620+SHP099 (daily), MRTX1257 (50 mg/kg, daily) or MRTX1257 +SHP099 (daily) at indicated time points. **H-I**, CD31, staining in *LSL-KRAS*$^{G12C}$- (H) and - *LSL-KRAS*$^{G12C}$-*Tp53*$^{R270H}$- (I) derived tumors after 3 days of treatment, as indicated.

**Supplementary Figure S7. ARS1620/SHP099 also evokes anti-tumor immune response in NSCLC GEMMs. A,** Pie charts showing immune cell populations in *LSL-KRAS*$^{G12C}$ tumors, treated as indicated for 6 days. **B,** Multiplex IF/IHC analysis of *LSL-KRAS*$^{G12C}$- and *LSL-KRAS*$^{G12C}$;*Tp53*$^{R270H}$ tumors, treated as indicated for 3 days, and stained with the indicated markers. **C,** Tumor-infiltrating immune cells from *LSL-KRAS*$^{G12C}$ tumors analyzed after 6 days of treatment.

**Table S1: Bliss index for effects of SHP099/ARS1620 combination on cancer cells**

| Proliferation | Bliss Independence Analysis | | | | |
|---|---|---|---|---|---|
| **Figure 1A** | **Ea** | **Eb** | **Bliss Independent** | **Observed** | **Synergistic** |
| **WT** | 45.55% | 5.56% | 48.58% | 48.48% | NO |
| **G12C** | 20.40% | 31.37% | 45.37% | 65.69% | YES |
| **G12D** | 6.28% | 6.39% | 12.27% | 6.01% | NO |
| **Q61R** | 11.92% | 5.21% | 16.51% | 17.08% | NO |
| **Figure 1B** | **Bliss Independence Analysis** | | | | |
| **2D** | **Ea** | **Eb** | **Bliss Independent** | **Observed** | **Synergistic** |
| **H358** | 53.15% | 91.47% | 96.00% | 100.00% | YES |
| **H1373** | 10.10% | 90.76% | 91.69% | 97.08% | YES |
| **H2122** | 8.59% | 55.97% | 59.75% | 93.60% | YES |
| **H1792** | 26.84% | 44.69% | 59.53% | 89.88% | YES |
| **H23** | 30.06% | 49.56% | 64.72% | 89.96% | YES |
| **CALU1** | 8.80% | 45.31% | 50.13% | 72.49% | YES |
| **H2030** | 3.27% | 14.66% | 17.45% | 35.59% | YES |
| **SW1573** | 16.24% | 4.61% | 20.10% | 20.53% | NO |
| **H460** | 2.88% | 1.79% | 4.61% | 0.64% | NO |
| **3D** | **Ea** | **Eb** | **Bliss Independent** | **Observed** | **Synergistic** |
| **H358** | 93.64% | 100.00% | 100.00% | 100.00% | NO |
| **H1373** | 34.90% | 82.94% | 88.90% | 99.09% | YES |
| **H2122** | 1.92% | 42.30% | 43.41% | 84.76% | YES |
| **H1792** | 44.12% | 53.76% | 74.16% | 75.55% | YES |
| **H23** | 70.91% | 74.44% | 92.56% | 92.92% | YES |
| **CALU1** | 88.18% | 100.00% | 100.00% | 100.00% | NO |
| **H2030** | 27.22% | 55.25% | 67.43% | 70.52% | YES |
| **SW1573** | 45.45% | 40.70% | 67.65% | 39.89% | NO |
| **H460** | 2.88% | 0.12% | 2.99% | 1.80% | NO |
| **Figure 1C** | **Bliss Independence Analysis** | | | | |
| | **Ea** | **Eb** | **Bliss Independent** | **Observed** | **Synergistic** |
| **MIAPACA 2D** | -9.64% | 5.93% | -3.13% | 90.74% | YES |
| **PANC0327 2D** | 36.80% | 13.63% | 45.41% | 39.53% | NO |
| **NY53 2D** | 3.76% | 5.03% | 8.61% | 33.86% | YES |
| **KCP** | 26.89% | 23.23% | 43.87% | 90.54% | YES |
| **KPC** | 30.50% | -5.12% | 26.95% | 26.07% | NO |

**Table S2. Blood counts after SHP099/ARS1620 treatment**

| | Mean±SD (n=4) | Ref. values |
|---|---|---|
| **WBC ($10^9$/L)** | 3.4±0.6 | 1.8 – 10.7 |
| **NE ($10^9$/L)** | 0.9±0.3 | 0.1 – 2.4 |
| **LY ($10^9$/L)** | 2.3±0.5 | 0.9 – 9.3 |
| **MO ($10^9$/L)** | 0.1±0.0 | 0.0 – 0.4 |
| **EO ($10^9$/L)** | 0.0±0.0 | 0.0 – 0.2 |
| **BA ($10^9$/L)** | 0.0±0.0 | 0.0 – 0.2 |
| **RBC ($10^{12}$/L)** | 9.3±0.8 | 6.36 – 9.42 |
| **Hb (g/dL)** | 12.8±1.2 | 11.0 – 15.1 |
| **HCT (%)** | 40.5±3.5 | 35.1 – 45.4 |
| **MCV (fL)** | 53.1±2.1 | 45.4 – 60.3 |
| **PLTS ($10^9$/L)** | 872±152 | 592 – 2972 |
| **MPV (fL)** | 5.0±0.5 | 5.0 – 20.0 |

*WBC: white blood cells; NE: neutrophils; LY: lymphocytes; MO: monocytes; EO: eosinophils; BA: basophils; RBC: red blood cells; Hb: hemoglobin; HCT: hematocrit; MCV: mean cell volume; PLTS: platelets; MPV: mean platelet volume*

**expected range from HemaVet 950FS (Drew Scientific)*

### Table S3. Primer, sg RNA and Fab Sequences

| G12C targeting primers | |
|---|---|
| **mKras INTRON 0 Forward (F1)** | GTCTTTCCCCAGCACAGTGC |
| **mKras WT site-specific Forward (W)** | ACTTGTGGTGGTTGGAGCTGG |
| **mKras G12D site-specific Forward (D)** | GCTTGTGGTGGTTGGAGCTGA |
| **mKras G12C site-specific Forward (C)** | ACTTGTCGTCGTTGGAGCTTG |
| **mKras INTRON 1 Reverse (R2)** | CCTTTACAAGCGCACGCAGACTGTAGAGC |

| RT-PCR:primers | |
|---|---|
| **ETV1 Forward** | CTTAGCCGTTCACTCCGCTAT |
| **ETV1 Reverse** | TCTGTCTTCAGCAGTGGACG |
| **ETV4 Forward** | GCCCATTTCATTGCCTGGAC |
| **ETV4 Reverse** | TACACGTAACGCTCACCAGC |
| **ETV5 Forward** | TAGAACCGGAAGAGGTTGCTC |
| **ETV5 Reverse** | TTATCCGGGAAAGCCATGGAG |
| **DUSP6 Forward** | TGGAGGAATTCGGCATCAAGT |
| **DUSP6 Reverse** | AGCAATGTACCAAGACACCACA |
| **EGF Forward** | CAGCTGTGTCATTGGATGTGC |
| **EGF Reverse** | ACGGTCACCAAAAAGGGACA |
| **FGF2 Forward** | AGCAGAAGAGAGAGGAGTTGTG |
| **FGF2 Reverse** | TCGTTTCAGTGCCACATACCA |
| **PDGFC Forward** | CTGGTTAAACGCTGTGGTGG |
| **PDGFC Reverse** | CTGACACCGGTCTTTGGTCT |
| **PDGFD Forward** | TGTGGCTGTGGAACTGTCAA |
| **PDGFD Reverse** | ATCGAGGTGGTCTTGAGCTG |
| **AXL Forward** | GACTATCTGCGCCAGGGAAA |
| **AXL Reverse** | TAAAACTTGGCCGGTCCTGG |
| **EGFR Forward** | GCGTCCGCAAGTGTAAGAAG |
| **EGFR Reverse** | TCCAGAGGAGGAGTATGTGTGA |
| **FGFR1 Forward** | CAGAGACCCACCTTCAAGCA |
| **FGFR1 Reverse** | AGCGGCTCATGAGAGAAGAC |
| **FGFR2 Forward** | GTGATGTCTGGTCCTTCGGG |
| **FGFR2 Reverse** | GAACGTTGGTCTCTGGGAGG |
| **FGFR3 Forward** | AGGAGCTCTTCAAGCTGCTG |
| **FGFR3 Reverse** | AGGTCCAGGTACTCGTCGG |
| **FGFR4 Forward** | CTCCCAGAGGCCTACCTTCA |
| **FGFR4 Reverse** | CACCAGAGGGGGAATAGGGT |
| **HER2 Forward** | CAGGAGTGCGTGGAGGAATG |
| **HER2 Reverse** | GGCCACACACTGGTCAGC |
| **HER3 Forward** | GTGGTGATGGGGAACCTTGA |
| **HER3 Reverse** | CGGAGGTTGGGCAATGGTAG |
| **IGF1R Forward** | CTTCGCTTCGTCATGGAGGG |
| **IGF1R Reverse** | CAGCTTGTTCTCCTCGCTGT |
| **MET Forward** | CTGAATCTGCAACTCCCCCT |
| **MET Reverse** | CCTTTAACTGCTTCAGGGTCAA |
| **PDGFRA Forward** | ACCACCCAGAGAAGCCAAAG |
| **PDGFRA Reverse** | GTATCAGCCTGCTTCATGTCC |
| **PDGFRB Forward** | AGCCCAATGAGGGTGACAAC |

| | |
|---|---|
| **PDGFRB Reverse** | TGACTTCATTCAGGGTGGAGC |
| **RET Forward** | GGATGGAGAGGCCAGACAAC |
| **RET Reverse** | GAGTCAGATGGAGTGGACGC |
| **RON Forward** | CCGCCACATTGACCCTTTTG |
| **RON Reverse** | AGCTGCACATAATGGTCCCC |
| **GAPDH Forward** | GAGTCAACGGATTTGGTCGT |
| **GAPDH Reverse** | TTGATTTTGGAGGGATCTCG |
| **Oligonucleotides used for cloning sgRNAs** | |
| **mKras G12D Forward** | CACCGAATGACTGAGTATAAGCTTG |
| **mKras G12D Reverse** | AAACCAAGCTTATACTCAGTCATTC |
| **mPtpn11 Forward** | CACCGAAAACTGCCATCGACTCCTC |
| **mPtpn11 Reverse** | AAACGAGGAGTCGATGGCAGTTTTC |
| **PTPN11 Forward** | CACCGGATTACTATGACCTGTATGG |
| **PTPN11 Reverse** | AAACCCATACAGGTCATAGTAATCC |
| **HDR ssDNA template** | |
| **G12C:** | CACACAAAGGTGAGTGTTAAAATATTGATAAAGTTTTTGATAATCTTGTGTGAGACATGT<br>TCTAATTTAGTTGTATTTTATTATTTTTATTGTAAGGCCTGCTGAAAATGACTGAGTATAA<br>ACTTGTCGTCGTTGGAGCTTGTGGCGTAGGCAAGAGCGCCTTGACGATACAGCTAATTC<br>AGAATCACTTTGTGGATGAG |
| **12C-ARS Fab sequence** | |
| **Light chain** | GATATCCAGATGACCCAGTCCCCGAGCTCCCTGTCCGCCTCTGTGGGCGATAGGGTCACC<br>ATCACCTGCCGTGCCAGTCAGTCCGTGTCCAGCGCTGTAGCCTGGTATCAACAGAAACCA<br>GGAAAAGCTCCGAAGCTTCTGATTTACTCGGCATCCAGCCTCTACTCTGGAGTCCCTTCTC<br>GCTTCTCTGGTAGCCGTTCCGGGACGGATTTCACTCTGACCATCAGCAGTCTGCAGCCGG<br>AAGACTTCGCAACTTATTACTGTCAGCAAGACTGGTACTTCCCGATCACGTTCGGACAGG<br>GTACCAAGGTGGAGATCAAACGAACTGTGGCTGCACCATCTGTCTTCATCTTCCCGCCAT<br>CTGATTCACAGTTGAAATCTGGAACTGCCTCTGTTGTGTGCCTGCTGAATAACTTCTATCC<br>CAGAGAGGCCAAAGTACAGTGGAAGGTGGATAACGCCCTCCAATCGGGTAACTCCCAG<br>GAGAGTGTCACAGAGCAGGACAGCAAGGACAGCACCTACAGCCTCAGCAGCACCCTGA<br>CGCTGAGCAAAGCAGACTACGAAAAACATAAAGTCTACGCCTGCGAAGTCACCCATCAG<br>GGCCTGAGCTCGCCCGTCACAAAGAGCTTCAACAGGGGAGAGTGT |
| **Heavy chain** | GAGGTTCAGCTGGTGGAGTCTGGCGGTGGCCTGGTGCAGCCAGGGGGCTCACTCCGTTT<br>GTCCTGTGCAGCTTCTGGCTTCACTTTCTCTTCTTATTATATACACTGGGTGCGTCAGGCC<br>CCGGGTAAGGGCCTGGAATGGGTTGCATCTATTTCTCCTTCTTCTGGCTCTACTTATTATG<br>CCGATAGCGTCAAGGGCCGTTTCACTATAAGCGCAGACACATCCAAAAACACAGCCTACC<br>TACAAATGAACAGCTTAAGAGCTGAGGACACTGCCGTCTATTATTGTGCTCGCTACGGTG<br>GTCGTTCTTACTGGCAGAAACAGGACTCTTACTTCTACCAGCATGGTTTGGACTACTGGG<br>GTCAAGGAACCCTGGTCACCGTCTCCTCGGCCTCCACCAAGGGTCCATCGGTCTTCCCCC<br>TGGCACCCTCCTCCAAGAGCACCTCTGGGGGCACAGCGGCCCTGGGCTGCCTGGTCAAG<br>GACTACTTCCCCGAACCGGTGACGGTGTCGTGGAACTCAGGCGCCCTGACCAGCGGCGT<br>GCACACCTTCCCGGCTGTCCTACAGTCCTCAGGACTCTACTCCCTCAGCAGCGTGGTGAC<br>CGTGCCCTCCAGCAGCTTGGGCACCCAGACCTACATCTGCAACGTGAATCACAAGCCCAG<br>CAACACCAAGGTCGACAAGAAAGTTGAGCCCAAATCTTGTGACAAAACTCACACA |

**Table S4: Antibodies for Flow Cytometry**

| Company | Cat# | Antigen | Chromophore |
|---|---|---|---|
| Fisher | 564279 | CD45 | BUV395 |
| Biolegend | 100336 | CD3e | BV421 |
| Fisher | 564933 | CD4 | BUV737 |
| Biolegend | 100740 | CD8a | BV570 |
| Biolegend | 103028 | CD44 | APC/Cy7 |
| Biolegend | 135231 | PD1 | BV711 |
| Biolegend | 106310 | CTLA4 | APC |
| Biolegend | 119425 | OX40 | PerCP/Cy5.5 |
| Biolegend | 119725 | Tim3 | BV785 |
| Biolegend | 126404 | Foxp3 | PE |
| Fisher | 50-112-4690 | Ki-67 | Alexa700 |
| Biolegend | 115540 | CD19 | BV650 |
| Biolegend | 101236 | CD11b | BV421 |
| Fisher | 564986 | CD11c | BUV737 |
| Biolegend | 128030 | Ly6C | BV570 |
| Biolegend | 108431 | Ly6G | BV711 |
| Biolegend | 123118 | F4/80 | APC/Cy7 |
| BioLegend | 108906 | Dx5 | FITC |
| BioLegend | 115534 | CD19 | PerCP5.5 |
| eBioscience | 12-1522-83 | CTLA4 | PE |
| BioLegend | 104536 | CD69 | PE-CF594 |
| eBioscience | 25-5870-82 | Tim3 | PE/Cy7 |
| eBioscience | 17-5773-82 | Foxp3 | APC |
| BioLegend | 100216 | CD3 | AF700 |
| BioLegend | 135224 | PD-1 | APC/Cy7 |
| BioLegend | 107616 | IA/IE | FITC |
| BioLegend | 127616 | Ly6G | PerCP5.5 |
| BioLegend | 100708 | CD8 | PE |
| BioLegend | 123146 | F4/80 | PE-CF594 |
| Biolegend | 103039 | CD44 | BV421 |
| BioLegend | 652411 | Ki-67 | BV421 |
| BioLegend | 124314 | PD-L1 | PECy7 |
| BioLegend | 104412 | CD62L | APC |
| BioLegend | 128026 | Ly6C | APC/Cy7 |
| BioLegend | 103140 | CD45 | BV605 |
| BioLegend | 100748 | CD8 | BV711 |
| BioLegend | 100453 | CD4 | BV785 |
| BioLegend | 101242 | CD11b | BV711 |
| BioLegend | 117336 | CD11c | BV785 |
